# COR27/28 Regulate the Evening Transcriptional Activity of the RVE8-LNK1/2 Circadian Complex

**DOI:** 10.1101/2022.05.16.492168

**Authors:** Maria L. Sorkin, Shin-Cheng Tzeng, Andrés Romanowski, Nikolai Kahle, Rebecca Bindbeutel, Andreas Hiltbrunner, Marcelo J. Yanovsky, Bradley S. Evans, Dmitri A. Nusinow

**Affiliations:** Donald Danforth Plant Science Center, St. Louis, MO, USA; Division of Biology and Biomedical Sciences, Washington University in St. Louis, St. Louis, MO, USA; Fundación Instituto Leloir, Instituto de Investigaciones Bioquímicas de Buenos Aires–Consejo Nacional de Investigaciones Científicas y Técnicas (CONICET), Buenos Aires, Argentina; Institute of Biology II, Faculty of Biology, University of Freiburg, Freiburg, Germany; Signalling Research Centres BIOSS and CIBSS, University of Freiburg, Freiburg, Germany

## Abstract

The timing of many molecular and physiological processes in plants occurs at a specific time of day. These daily rhythms are driven by the circadian clock, a master timekeeper that uses daylength and temperature to maintain rhythms of approximately 24 hours in various clock-regulated phenotypes. The circadian MYB-like transcription factor REVEILLE 8 (RVE8) interacts with its transcriptional coactivators NIGHT LIGHT INDUCIBLE AND CLOCK REGULATED 1 (LNK1) and LNK2 to promote the expression of evening-phased clock genes and cold tolerance factors. While genetic approaches have commonly been used to discover new connections within the clock and between other pathways, here we use affinity purification coupled with mass spectrometry to discover time-of-day-specific protein interactors of the RVE8-LNK1/2 complex. Among the interactors of RVE8/LNK1/LNK2 were COLD REGULATED GENE 27 (COR27) and COR28, which were coprecipitated in an evening-specific manner. In addition to COR27/28, we found an enrichment of temperature-related interactors that led us to establish a novel role for LNK1/2 in temperature entrainment of the clock. We established that RVE8, LNK1, and either COR27 or COR28 form a tripartite complex in yeast and that the effect of this interaction *in planta* serves to antagonize transcriptional activation of RVE8 target genes through mediating RVE8 protein degradation in the evening. Together, these results illustrate how a proteomic approach identified time-of-day-specific protein interactions and a novel RVE8-LNK-COR protein complex that implicates a new regulatory mechanism for circadian and temperature signaling pathways.

## Introduction

Daily and seasonal patterns in daylength and temperature cycles are two of the most dependable environmental cues an organism experiences. As such, lifeforms in every kingdom have evolved a mechanism to anticipate and synchronize their biology with the earth’s predictable 24-hour and 365-day cycles (Ouyang et al., 1998; Rosbash, 2009; Edgar et al., 2012). This mechanism is called the circadian clock, which in plants consists of approximately 20-30 genes that participate in transcription-translation feedback loops to produce rhythms with a period of about 24 hours (Creux and Harmer, 2019). These core oscillator genes respond to the environment by producing a physiological response appropriate for a particular time of day or year (Webb et al., 2019). In plants, the clock regulates a variety of phenotypic outputs, including the transition from vegetative to reproductive growth, biotic defense responses, and protection from abiotic stressors such as extreme warm or cold temperature (Greenham and Mcclung, 2015).

Identification of circadian-associated genes has been critical in understanding the generation of biological rhythms. Core oscillator components often exhibit rhythmic gene expression with a period of ~24 hours and a set phase—or time of peak and trough expression. For example, two of the first genes to be defined as core oscillator components in the model plant *Arabidopsis thaliana* (Arabidopsis) are the morning-phased MYB-like transcription factors *CIRCADIAN CLOCK ASSOCIATED 1* (*CCA1*) and *LATE ELONGATED HYPOCOTYL (LHY*) (Schaffer et al., 1998; Wang and Tobin, 1998; Green and Tobin, 1999). These genes are highly expressed at dawn and repress the expression of the afternoon- and evening-phased *PSEUDO RESPONSE REGULATOR* genes *PRR1 TIMING OF CAB EXPRESSION 1* (*TOC1*), *PRR5, PRR7*, and *PRR9* (Alabadí et al., 2001; Farré et al., 2005; Kamioka et al., 2016). The *PRRs* reciprocally repress *CCA1*/*LHY*, completing one of the negative feedback loops that define the clock. In the evening, *EARLY FLOWERING 3* (*ELF3*), *ELF4*, and *LUX ARRHYTHMO* (*LUX*) interact in the nucleus to form a tripartite protein complex called the evening complex, which represses *PRR9*, *CCA1/LHY*, and other clock and growth-promoting factors (Dixon et al., 2011; Nusinow et al., 2011; Chow et al., 2012; Herrero et al., 2012). As we discover new connections within and between the clock, we enhance our understanding of this important system.

In this study, we used affinity purification coupled with mass spectrometry (APMS) to identify protein-protein interactions associated with the REVEILLE 8 (RVE8)-NIGHT LIGHT-INDUCIBLE AND CLOCK-REGULATED 1 (LNK1)/LNK2 circadian transcriptional complex. The RVEs are an 8-member family of CCA1/LHY-like transcription factors of which some members interact with the LNK proteins to coregulate target gene expression (Rawat et al., 2011; Rugnone et al., 2013; Xie et al., 2014; Pérez-García et al., 2015; Gray et al., 2017). In the late morning, the RVE8-LNK1/2 transcriptional complex activates the expression of evening-expressed clock genes such as *TOC1* and *PRR5* via recruitment of the basal transcriptional machinery to these and other *RVE8* target promoters (Xie et al., 2014; Ma et al., 2018). Conversely, *LNK1/2* are also known to act as corepressors of other *RVE8* targets, such as the anthocyanin structural gene *UDP-GLUCOSE:FLAVONOID 3-O-GLUCOSYLTRANSFERASE* (*UF3GT*) (Pérez-García et al., 2015). Additionally, LNK1/2 interact with another transcription factor, MYB3, as corepressors to inhibit the expression of the phenylpropanoid biosynthesis gene *C4H* (Zhou et al., 2017). The mechanism behind the corepressive function of the LNKs and how they switch between an activating and a repressive role is unknown.

LNK1/2 bind to RVE8 and MYB3 via two conserved arginine/asparagine-containing motifs called R1/R2 located in the LNK C-terminus (Xie et al., 2014; Zhou et al., 2017). Additionally, the Extra N-terminal Tail (ENT) domain present in LNK1/2 but not LNK3/4 is required for their repressive activity with MYB3 (Zhou et al., 2017). The LNKs have no other known functional protein domains apart from these regions. RVE8 and the other RVEs are characterized by the presence of a LHY-/CCA1-LIKE (LCL) domain, which can directly bind the LNKs, presumably at the C-terminus (de Leone et al., 2018; Ma et al., 2018). RVE8 target gene promoters frequently contain the canonical *CCA1/LHY-binding* motif called the evening element (EE) as well as G-box-like and morning element (ME)-like motifs (Hsu et al., 2013a).

In addition to regulating circadian rhythms, *RVE4/8* regulate thermotolerance under both high and low temperatures (Li et al., 2019; Kidokoro et al., 2021). After exposure to heat shock, RVE4/8 upregulate the expression of *ETHYLENE RESPONSIVE FACTOR 53* (*ERF53*) and *ERF54*, boosting the plant’s heat shock tolerance (Li et al., 2019). In another study, the authors found that *RVE4/8* also appear to promote freezing tolerance via activation of *DEHYDRATION-RESPONSIVE ELEMENT BINDING PROTEIN 1A* (*DREB1A*, also referred to as *C-REPEAT BINDING FACTOR 3, CBF3*) when grown at 4°C (Kidokoro et al., 2021). A corresponding association between temperature and the LNKs has not been well studied, although EC-mediated induction of *LNK1* expression under warm nights suggests a role for the LNKs in temperature responses (Mizuno et al., 2014).

Our proteomic approach presented here establishes novel protein interactions with the RVE8-LNK1/2 transcriptional complex at ZT5 and ZT9. Although these clock bait proteins exhibit peak mRNA expression in the early morning hours, we found that LNK1 and RVE8 interact with more protein partners at the later ZT9 timepoint than at ZT5. Temperature response related GO terms were significantly enriched among the coprecipitated proteins, prompting us to explore and establish a role for LNK1/2 in temperature entrainment of the clock. Among the temperature-related coprecipitated proteins were COLD REGULATED GENE 27 (COR27) and COR28, which only coprecipitated with RVE8/LNK1/LNK2 at ZT9. Furthermore, we found that the CORs interact with RVE8 and LNK1 in a tripartite complex in a yeast 3-hybrid system. By performing APMS using 35S::YFP-COR27 and 35S::GFP-COR28, we validated the interaction with LNK1, LNK2, and RVE8, and identified additional novel interactions between the CORs and RVE5, RVE6, and several light signaling proteins. Further investigation into the role of the RVE8-LNK1/2-COR27/28 interaction suggested that the CORs antagonize activation of RVE8 target genes via regulation of RVE8 protein stability in the evening. Thus, by taking a proteomic approach to study a core circadian transcriptional complex, we identified a novel, evening-phased RVE8-LNK-COR protein complex that presents a new regulatory mechanism for circadian and temperature signaling pathways.

## Results

### Characterization of affinity-tagged lines

To identify new interactions with known clock proteins, we created endogenous promoter-driven, 3x-FLAG-6x-His C-terminal (HFC) affinity-tagged versions of RVE8, LNK1, and LNK2. RVE8-HFC was transformed into the *rve8-1 CCR2::LUC* mutant background while LNK1-HFC and LNK2-HFC were introduced into *lnk1/2/3/4* quadruple mutant (*lnkQ*) (de Leone et al., 2018) *CCA1::LUC*. By transforming our tagged LNKs into the *lnkQ* background, we could eliminate co-precipitating interactors that could be formed through a complex between our tagged LNKs and the endogenous LNKs. To ensure the tagged versions of our proteins of interest functioned similarly to their native counterparts, we selected T3 homozygous lines that rescued the long period mutant phenotype of *rve8-1* or *lnkQ* mutants (Rawat et al., 2011; Xie et al., 2014) (**Fig. 1A-C**). LNK1-HFC/LNK2-HFC did not fully restore the circadian period back to wild-type levels, but the lengthened period is consistent with the absence of the other three LNKs after the introduction of the tagged LNK into the *lnkQ* quadruple mutant (Xie et al., 2014; de Leone et al., 2018). We also determined that the HFC-tagged proteins exhibit rhythmic protein abundance patterns under 12 hr light: 12 hr dark (LD) conditions, as would be expected for these proteins (**Fig. 1D-G**). While mRNA expression for *RVE8, LNK1*, and *LNK2* peaks at ZT1, ZT5, and ZT2, respectively, peak protein abundance occurred at ZT6, ZT9, and ZT6—about 4-5 hours after peak mRNA expression (Mockler et al., 2007) (**Fig. 1D-G, Fig. S1**). This lag in protein abundance after transcription is consistent with previously reported data showing a peak in RVE8-HA abundance three to six hours after dawn (Rawat et al., 2011). These experiments demonstrate that our affinity-tagged clock proteins behaved similarly to the native protein and are functional, making them ideal tools for capturing relevant protein interactions.

**Figure 1.**
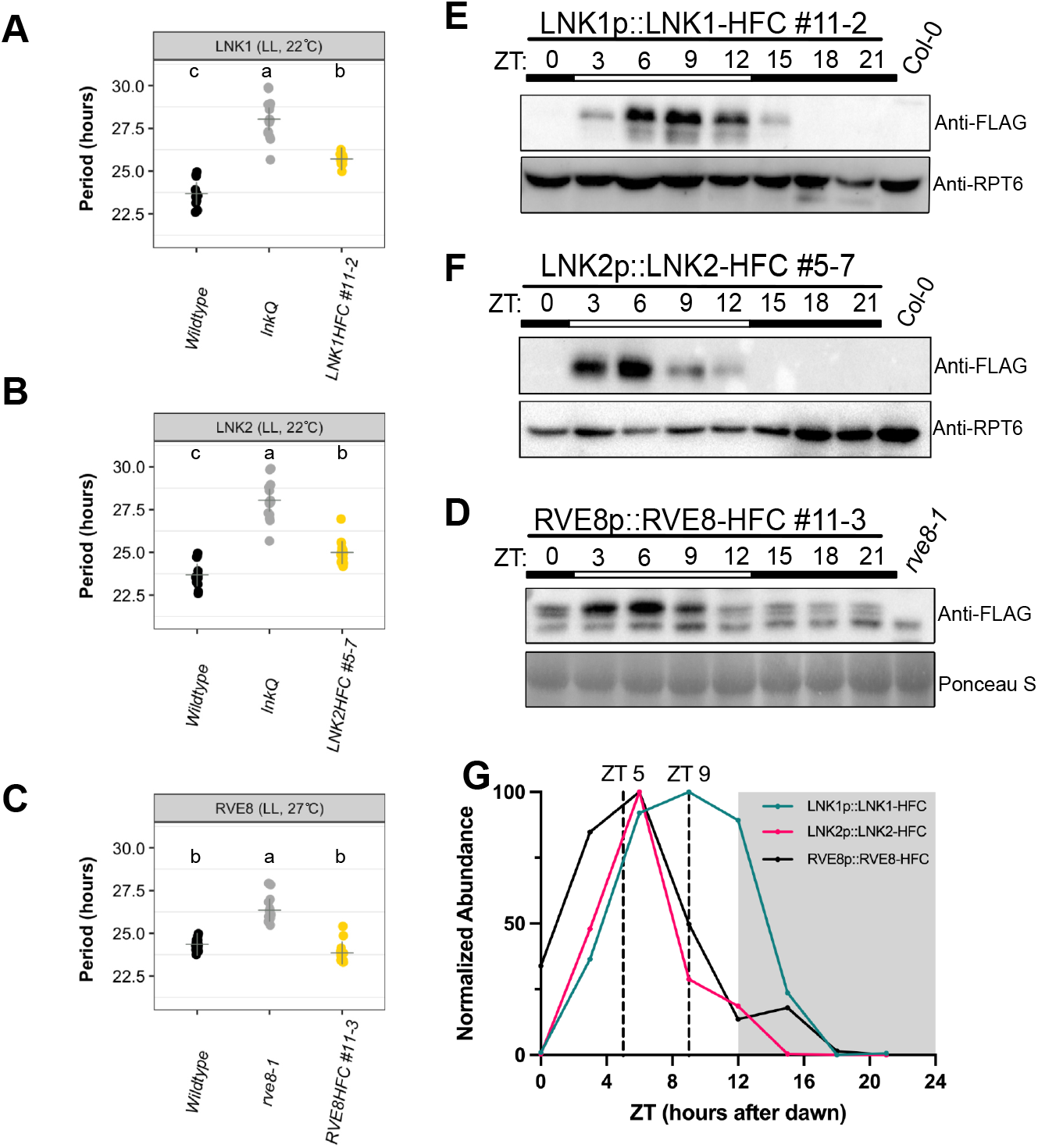
Characterization of affinity-tagged lines used for APMS. (A-C) Circadian luciferase reporter period analysis of selected T3 homozygous lines expressing (A) LNK1-HFC, (B) LNK2-HFC, or (C) RVE8-HFC in their respective mutant backgrounds (*rve8-l* or *lnkQ*). Each point represents the circadian period of an individual plant and the + symbol shows the average period for that genotype. Letters correspond to significantly different periods as determined by ANOVA with a Tukey’s post-hoc test. LNK1 and LNK2 luciferase assays were performed together and include the same wildtype and *InkQ* data. Environmental conditions during imaging are included at the top of the plot (LL = constant light). (D-F) Time course Western blots showing cyclic protein abundance patterns of 10-day-old affinity tagged lines under 12 hr light: 12 hr dark 22 °C conditions. Affinity tagged lines are detected with anti-FLAG antibody. RPT5 or Ponceau S staining was used to show loading. Col-0 CCA1::LUC (Col-0) or *rve8-l* CCR2::LUC (*rve8-l*) were used as negative controls. White and black bars indicate lights-on and lights-off, respectively (D) 24-hour protein expression patterns of affinity tagged lines normalized to Ponceau S or RPT5 quantified by densitometry of Western blots shown in D-F. Vertical dotted lines indicate time of tissue collection for APMS. White and grey shading indicates lights-on and lights-off, respectively. Western blots and luciferase reporter assays were repeated at least 2 times. ZT= Zeitgeber Time.

### Affinity purification-mass spectrometry (APMS) identifies novel time-of-day-specific interacting partners for RVE8, LNK1, and LNK2

We selected two timepoints for APMS based on the protein abundance patterns for RVE8-HFC, LNK1-HFC, and LNK2-HFC **(Fig. 1G**). RVE8-HFC and LNK2-HFC exhibited the highest protein abundance between ZT3 and ZT6, while LNK1-HFC protein was highest between ZT6 and ZT9. Considering this, we chose to examine protein-protein interactions at ZT5 and ZT9.

We identified a total of 392 proteins that coprecipitated with either RVE8-HFC, LNK1-HFC, or LNK2-HFC at ZT5 or ZT9 but did not coprecipitate in our GFP-HFC nor Col-0 negative controls (**Fig. 2A, Dataset S1**). Consistent with the time of peak LNK1-HFC and LNK2-HFC protein abundance (ZT9 and ZT5, respectively; **Fig. 1G**), we saw higher total spectra mapping to LNK1-HFC at ZT9 (621) and LNK2-HFC at ZT5 (497) compared to the other timepoint (**Tables 1 and 2**). Similarly, the number of coprecipitated proteins was greatest at ZT9 for LNK1-HFC and at ZT5 for LNK2-HFC (**Fig. 2B-C, Dataset S1**). Total spectra mapping to the bait protein RVE8-HFC were similar between the two timepoints (**Tables 1 and 2**). Despite the similarity in RVE8-HFC total spectra between timepoints, we precipitated more ZT9-specific interactors than ZT5-specific interactors with RVE8-HFC (**Fig. 2D**). Overall, we identified more RVE8/LNK1/LNK2-binding partners at ZT9 (364) versus the earlier timepoint of ZT5 (281) (**Fig. 2A**) and found that 111 out of 392 (28.3%) total proteins coprecipitated were ZT9-specific; these proteins were not coprecipitated in any APMS experiment performed at ZT5. In summary, the enrichment of coprecipitated proteins at ZT9 suggests an important post-translational role for the RVE8-LNK1/2 complex in the evening.

**Figure 2.**
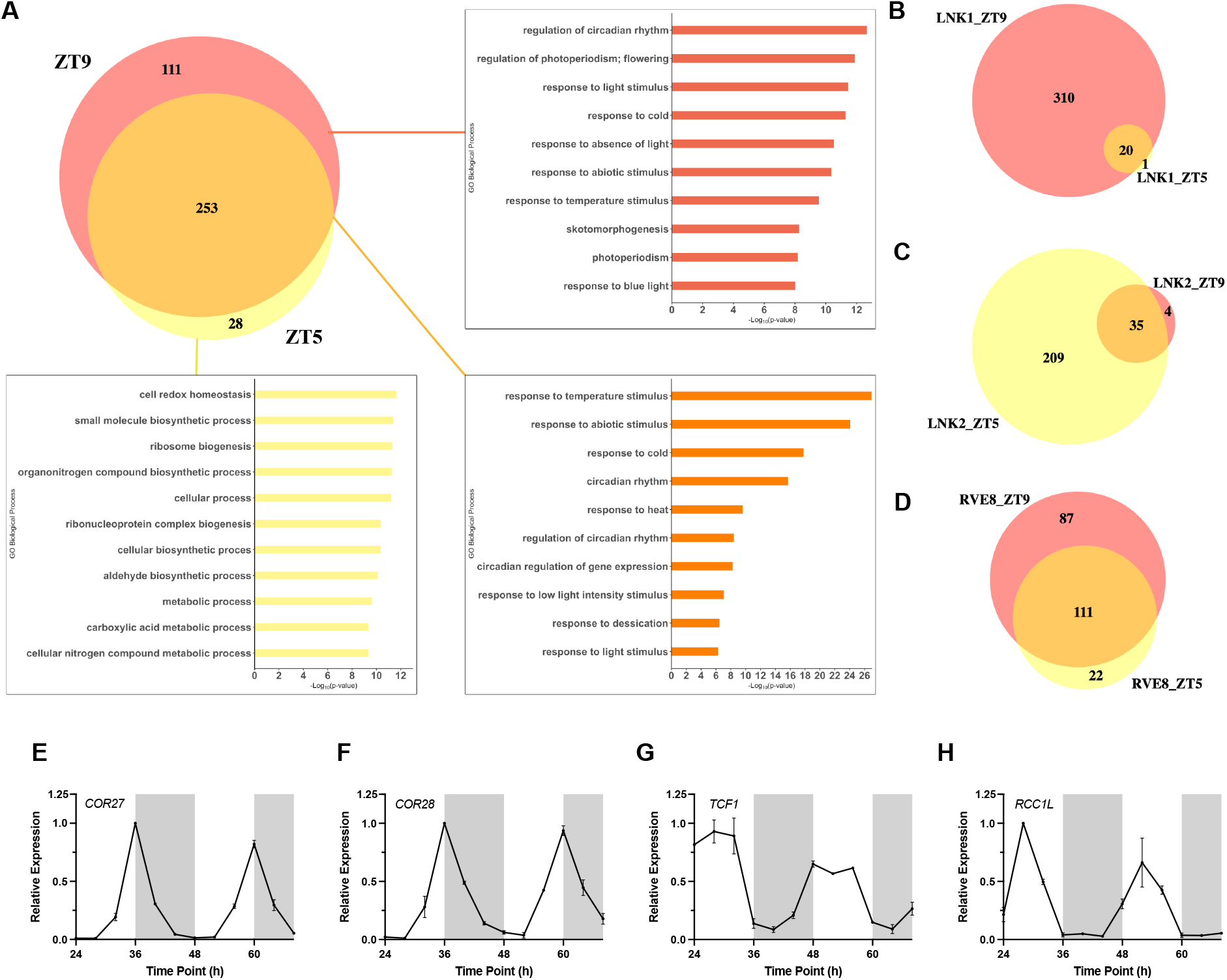
Analysis of proteins coprecipitated with RVE8/LNK1/LNK2-HFC by time-of-day affinity purification-mass spectrometry. (A) Venn diagram showing number of proteins coprecipitated with RVE8/LNK1/LNK2 at ZT5, ZT9, or at both timepoints. Corresponding bar charts show enriched GO biological process terms with -Log10_(p-value)_. (B-D) Venn diagrams of coprecipitated proteins at ZT5 and ZT9 separated by bait protein (B, LNK1-HFC, C, LNK2-HFC, or D, RVE8-HFC). (E-H) mRNA expression profiles in constant light of four cold-response proteins identified as RVE8/LNK1/LNK2 interactors. RNA-seq data for E-H taken from Romanowski et al. (2020) *The Plant Journal*. ZT= Zeitgeber Time

**Table 1.**
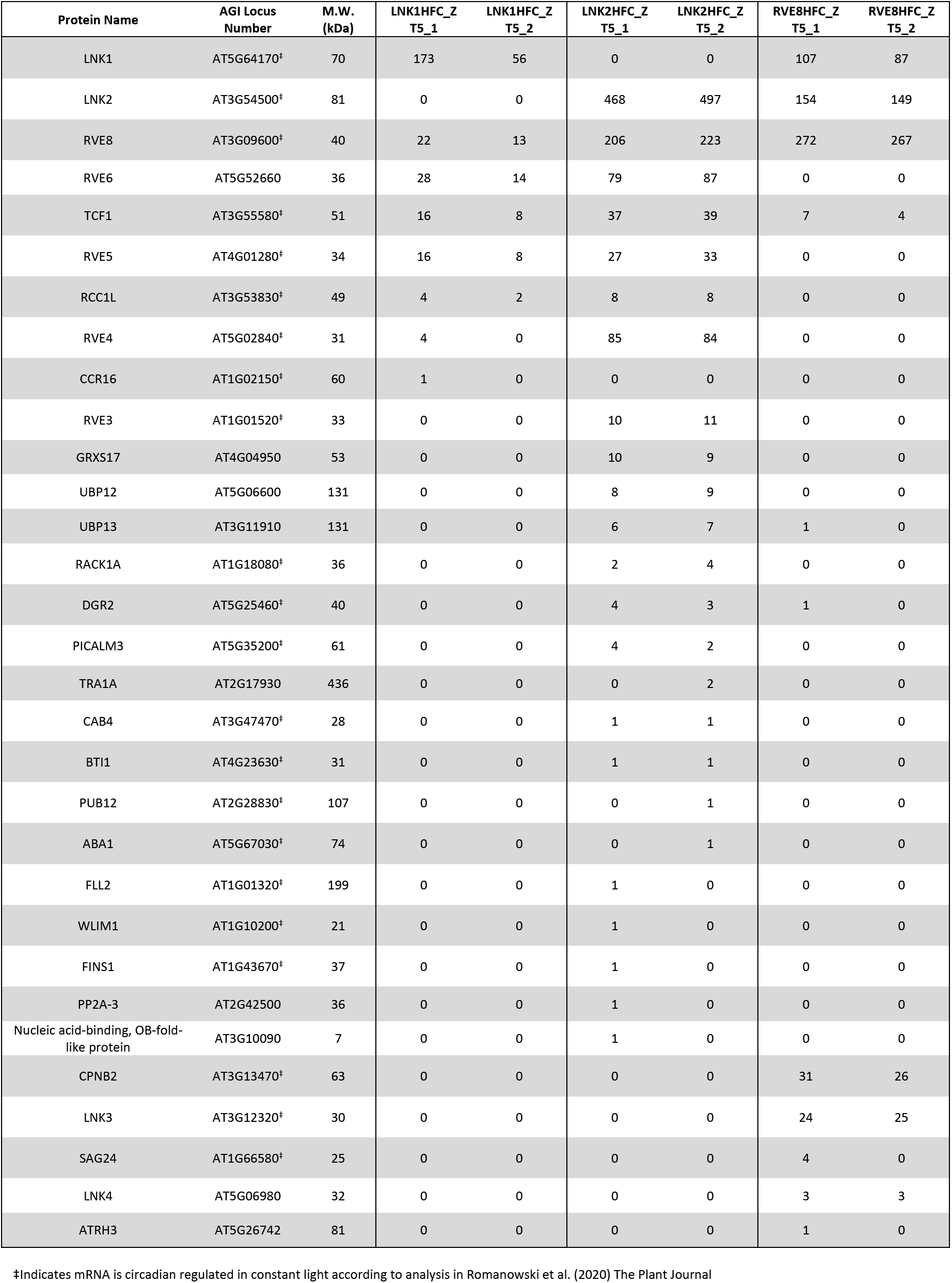
Proteins coprecipitated with RVE8/LNK1/LNK2-HFC at ZT5. Total spectra for a given coprecipitated protein is shown for each independent ZT5 sample. The curated table excludes coprecipitated proteins that were identified in the GFP-HFC or Col-0 negative control APMS experiments, see Dataset S1 for all identifications.

**Table 2.** Identified proteins coprecipitated with RVE8/LNK1/LNK2-HFC at ZT9. Total spectra for a given coprecipitated protein is shown for each independent ZT9 sample. The curated table excludes coprecipitated proteins that were identified in the GFP-HFC or Col-0 negative control APMS experiments, see Dataset S1 for all identifications.

| Protein Name | AGI Locus Number | M.W. (kDa) | LNK1HFC_Z T9_1 | LNK1HFC_Z T9_2 | LNK2HFC_Z T9_1 | LNK2HFC_Z T9_2 | RVE8HFC_Z T9_1 | RVE8HFC_Z T9_2 |
| --- | --- | --- | --- | --- | --- | --- | --- | --- |
| LNK1 | AT5G64170<sup>‡</sup> | 70 | 435 | 621 | 0 | 0 | 66 | 90 |
| LNK2 | AT3G54500<sup>‡</sup> | 81 | 0 | 0 | 140 | 151 | 109 | 137 |
| RVE8 | AT3G09600<sup>‡</sup> | 40 | 71 | 101 | 33 | 52 | 285 | 317 |
| RVE6 | AT5G52660 | 36 | 86 | 113 | 21 | 22 | 4 | 3 |
| RVE5 | AT4G01280<sup>‡</sup> | 34 | 32 | 51 | 12 | 12 | 0 | 0 |
| TCF1 | AT3G55580<sup>‡</sup> | 51 | 32 | 49 | 20 | 30 | 13 | 13 |
| RVE4 | AT5G02840<sup>‡</sup> | 31 | 31 | 43 | 0 | 7 | 0 | 0 |
| RCC1L | AT3G53830<sup>‡</sup> | 49 | 20 | 25 | 3 | 10 | 0 | 0 |
| COR28 | AT4G33980<sup>‡</sup> | 26 | 11 | 15 | 6 | 6 | 26 | 29 |
| RVE3 | AT1G01520<sup>‡</sup> | 33 | 8 | 10 | 0 | 0 | 0 | 0 |
| CCR16 | AT1G02150<sup>‡</sup> | 60 | 3 | 8 | 0 | 0 | 0 | 1 |
| TRA1A | AT2G17930 | 436 | 3 | 7 | 0 | 0 | 0 | 0 |
| DGR2 | AT5G25460<sup>‡</sup> | 40 | 3 | 4 | 0 | 0 | 2 | 5 |
| ENTH/ANTH/VHS superfamily protein | AT5G35200<sup>‡</sup> | 61 | 2 | 3 | 0 | 0 | 0 | 0 |
| FLL2 | AT1G01320<sup>‡</sup> | 199 | 0 | 3 | 0 | 0 | 0 | 1 |
| Nucleic acid-binding, OB-fold-like protein | AT2G40660<sup>‡</sup> | 42 | 3 | 2 | 0 | 0 | 0 | 0 |
| UBP12 | AT5G06600 | 131 | 0 | 1 | 0 | 0 | 0 | 0 |
| PHOT2 | AT5G58140 | 102 | 1 | 2 | 0 | 0 | 0 | 0 |
| MLK4 | AT3G13670<sup>‡</sup> | 79 | 0 | 2 | 0 | 0 | 6 | 6 |
| UBP13 | AT3G11910 | 131 | 0 | 2 | 0 | 0 | 1 | 4 |
| COR27 | AT5G42900 | 27 | 2 | 1 | 0 | 1 | 4 | 7 |
| WLIM1 | AT1G10200<sup>‡</sup> | 21 | 1 | 1 | 0 | 0 | 1 | 2 |
| CAB4 | AT3G47470<sup>‡</sup> | 28 | 1 | 1 | 0 | 0 | 1 | 1 |
| MLK2 | AT3G03940 | 78 | 0 | 0 | 1 | 1 | 6 | 7 |
| GRXS17 | AT4G04950 | 53 | 0 | 0 | 0 | 0 | 1 | 2 |
| CPNB2 | AT3G13470<sup>‡</sup> | 63 | 0 | 0 | 0 | 0 | 35 | 41 |
| LNK3 | AT3G12320<sup>‡</sup> | 31 | 0 | 0 | 0 | 0 | 18 | 23 |
| LNK4 | AT5G06980 | 32 | 0 | 0 | 0 | 0 | 2 | 6 |
| COP1 | AT2G32950<sup>‡</sup> | 76 | 0 | 0 | 0 | 0 | 3 | 4 |
| MLK1 | AT5G18190 | 77 | 0 | 0 | 0 | 0 | 3 | 4 |
| SPA1 | AT2G46340<sup>‡</sup> | 115 | 0 | 0 | 0 | 0 | 1 | 4 |
| MLK3 | AT2G25760 | 76 | 0 | 0 | 0 | 0 | 3 | 0 |

<sup>‡</sup>Indicates mRNA is circadian regulated in constant light according to analysis in Romanowski et al. (2020) *The Plant Journal*

We used gene ontology (GO) analysis to categorize coprecipitated proteins at ZT5, ZT9, and ZT5/9 (**Fig. 2A**). Proteins coprecipitated at ZT5 only were mostly assigned GO biological process terms associated with homeostasis and general metabolism while proteins found at ZT9 only or ZT5/ZT9 fell into relevant categories such as ‘regulation of circadian rhythm’, ‘response to light stimulus’, and ‘photoperiodism’ (**Fig. 2A**). We also noted that GO terms associated with temperature response were enriched in our interactor dataset (‘response to cold’, ‘response to temperature stimulus’, and ‘response to heat’) (**Fig. 2A**). This analysis suggested that we identified biologically relevant interacting partners involved in circadian rhythms in our APMS experiments and that there is an enrichment of temperature-related factors among these interactors. We also cross-referenced our lists of coprecipitated proteins with known cycling genes (Romanowski et al., 2020) and found that 71.0% of ZT5 and 71.1% of ZT9 proteins exhibited cyclic mRNA expression (**Dataset S1**), demonstrating that our bait circadian clock proteins mostly interacted with proteins whose expression also cycles.

Among the top interactors for LNK1-HFC, LNK2-HFC, and RVE8-HFC were four cold-response proteins: COLD REGULATED GENE 27 (COR27), COR28, and two regulator of chromosome condensation family proteins, TOLERANT TO CHILLING/FREEZING 1 (TCF1), and a homolog of TCF1 that we named REGULATOR OF CHROMOSOME CONDENSATION 1-LIKE (*RCC1L*, AT3G53830) (**Tables 1-2**). We characterized these as high-priority interactors based on their subcellular localization prediction and mRNA expression patterns; all four proteins are predicted to be nuclear localized according to the SUBACon subcellular localization consensus algorithm (Hooper et al., 2014), which stands in agreement with being interactors of the nuclear-localized RVE8/LNK1/LNK2 proteins; additionally, the mRNA expression for these genes is rhythmic under constant light conditions, suggesting circadian regulation of their expression (**Fig. 2E-H**). TCF1 and RCC1L were coprecipitated with RVE8/LNK1/LNK2 at both ZT5 and ZT9 while COR27/28 were ZT9-specific interactors **(Tables 1-2**).

*TCF1* and *RCC1L* are homologs of the regulator of chromosome condensation (RCC) family protein, RCC1 (Ji et al., 2015) and share 49.7% identity in an amino acid alignment (**Fig. S2**). RCC1 is a highly conserved guanine nucleotide exchange factor (GEF) for the GTP-binding protein RAN and is involved in nucleocytoplasmic export along with regulation of the cell cycle via chromosome condensation during mitosis (Ren et al., 2020). While there are no previous publications characterizing *RCC1L*, its sister gene *TCF1* is a known negative regulator of cold tolerance in Arabidopsis via the lignin biosynthesis pathway (Ji et al., 2015). *RCC1L* expression is downregulated upon cold treatment (**Table S1**), but no formal studies have been made into its role in cold tolerance nor chromatin biology.

COR27/28 have no known protein domains and are repressors of genes involved in cold tolerance, circadian rhythms and photomorphogenesis (Li et al., 2016; Wang et al., 2017; Kahle et al., 2020; Li et al., 2020; Zhu et al., 2020). Notably, COR27/28 repress the same clock and cold tolerance genes that are activated by RVE8; *PRR5, TOC1*, and *DREB1A* are repressed by the CORs and activated by RVE8 (Rawat et al., 2011; Kidokoro et al., 2021). Null or knock-down mutants of *cor27/cor28* exhibit a long period mutant phenotype, similar to that observed for *lnk* and *rve8* mutants (Rawat et al., 2011; Rugnone et al., 2013; Li et al., 2016). As the CORs do not contain a known DNA-binding domain, it is not understood how, mechanistically, these factors alter transcription.

Among the 111 evening-specific interactors were COR27, COR28, CONSTITUTIVELY PHOTOMORPHOGENIC 1 (COP1), and SUPPRESSOR OF PHYA-105 (SPA1) (**Table 2**). COP1 and SPA1 were RVE8-HFC-specific interactors while COR27/28 coprecipitated at ZT9 with LNK1/LNK2/RVE8-HFC. We hypothesized that this time-of-day-specific coprecipitation could be explained by the relative abundance of these proteins at ZT5 versus ZT9 due to diurnal changes in gene expression over the course of the day. To investigate this hypothesis, we overlayed the LD mRNA expression patterns of these ZT9-specific interactors on top of the protein abundance levels of RVE8-HFC, LNK1-HFC, and LNK2-HFC that were determined by time course Western blots shown in **Figure 1D-F** (**Fig. S3**). There is very little overlap in expression between the *CORs* and *RVE8/LNK1/LNK2* at ZT5 (**Fig. S3)**, indicating that COR27/28 may have only coprecipitated at ZT9 due to increased expression at that timepoint. In contrast, there was not a clear time-of-day distinction in expression overlap between COP1/SPA1 and the clock bait proteins, suggesting the ZT9-specific interaction between COP1/SPA1 and RVE8-HFC is possibly due to a factor other than expression level, such as recruitment through other proteins (such as COR27 or COR28) (**Fig. S3**).

### COR27 and COR28 interact with circadian and light signaling proteins

To better understand the role of COR27/28 at the protein level, we performed APMS using 35S::YFP-COR27 and 35S::GFP-COR28 lines (Li et al., 2016) collected at ZT9. Through this experiment, we validated the interactions between the CORs and RVE8/LNK1/LNK2 and additionally coprecipitated RVE5 and RVE6, further supporting the connection between COR27/28 and the RVE/LNK proteins (**Table 3, Dataset S1**). Previous studies have shown an interaction between COR27/28 and PHYTOCHROME B (PHYB), COP1, and SPA1 (Kahle et al., 2020; Li et al., 2020; Zhu et al., 2020). Our affinity purification captured these known interactions and additionally identified PHYD and SPA2/3/4, supporting the previously demonstrated role for the CORs in photomorphogenesis (**Table 3**) (Kahle et al., 2020; Li et al., 2020; Zhu et al., 2020). TCF1, one of the cold-tolerance proteins (Ji et al., 2015) to coprecipitate with RVE8/LNK1/LNK2, was also captured with COR27 (**Table 3**), which further implicates the CORs in freezing tolerance. In total, we identified 268 proteins that coprecipitated with YFP-COR27 or GFP-COR28 (**Dataset S1**). Of these, we found 58.9% exhibited circadian-regulated mRNA (Romanowski et al., 2020) (**Dataset S1**). Together, the COR27/28 APMS provides strong evidence that these proteins are important factors in circadian and light signaling networks.

**Table 3.** Identified proteins coprecipitated with YFP-COR27/GFP-COR28 at ZT9. Total spectra for a given coprecipitated protein is shown for each independent ZT9 sample. The curated table excludes coprecipitated proteins that were identified in the GFP-HFC or Col-0 negative control APMS experiments, see Dataset S1 for all identifications.

| Protein Name | AGI Locus Number | M.W. (kDa) | YFP-COR27_ZT9_1 | YFP-COR27_ZT9_2 | YFP-COR27_ZT9_3 | YFP-COR27_ZT9_4 | GFP-COR28_ZT9_1 | GFP-COR28_ZT9_2 | GFP-COR28_ZT9_3 | GFP-COR28_ZT9_4 |
| --- | --- | --- | --- | --- | --- | --- | --- | --- | --- | --- |
| COR27 | AT5G42900 | 27 | 89 | 85 | 95 | 80 | 0 | 0 | 0 | 0 |
| COR28 | AT4G33980<sup>‡</sup> | 26 | 0 | 0 | 0 | 0 | 22 | 18 | 8 | 10 |
| COP1 | AT2G32950<sup>‡</sup> | 76 | 16 | 12 | 19 | 16 | 6 | 6 | 1 | 2 |
| SPA1 | AT2G46340<sup>‡</sup> | 115 | 16 | 12 | 16 | 13 | 4 | 6 | 1 | 0 |
| MLK4 | AT3G13670<sup>‡</sup> | 79 | 16 | 11 | 15 | 12 | 0 | 0 | 0 | 0 |
| MLK2 | AT3G03940 | 78 | 13 | 12 | 15 | 12 | 0 | 0 | 0 | 0 |
| PHYD | AT4G16250 | 129 | 8 | 7 | 13 | 11 | 0 | 0 | 0 | 0 |
| MLK1 | AT5G18190 | 77 | 10 | 10 | 12 | 10 | 0 | 0 | 0 | 0 |
| SPA4 | AT1G53090 | 89 | 8 | 4 | 11 | 7 | 0 | 0 | 0 | 0 |
| SPA2 | AT4G11110 | 115 | 8 | 6 | 10 | 6 | 0 | 1 | 0 | 0 |
| SF1 | AT5G51300 | 87 | 7 | 12 | 7 | 5 | 0 | 0 | 0 | 0 |
| RVE8 | AT3G09600<sup>‡</sup> | 40 | 7 | 5 | 7 | 6 | 3 | 4 | 0 | 3 |
| LNK2 | AT3G54500<sup>‡</sup> | 81 | 6 | 5 | 6 | 3 | 0 | 1 | 0 | 0 |
| MLK3 | AT2G25760 | 76 | 5 | 6 | 6 | 8 | 0 | 0 | 0 | 0 |
| LNK1 | AT5G64170<sup>‡</sup> | 70 | 4 | 4 | 5 | 4 | 4 | 3 | 1 | 0 |
| SPA3 | AT3G15354<sup>‡</sup> | 93 | 4 | 4 | 4 | 4 | 0 | 0 | 0 | 0 |
| RVE6 | AT5G52660 | 36 | 3 | 3 | 4 | 1 | 1 | 0 | 0 | 0 |
| RVE5 | AT4G01280<sup>‡</sup> | 34 | 1 | 1 | 2 | 0 | 1 | 0 | 0 | 0 |
| CCR2 | AT2G21660<sup>‡</sup> | 17 | 1 | 0 | 1 | 1 | 0 | 0 | 0 | 0 |
| TCF1 | AT3G55580<sup>‡</sup> | 51 | 0 | 0 | 1 | 0 | 0 | 0 | 0 | 0 |
| PHYE | AT4G18130<sup>‡</sup> | 123 | 1 | 0 | 0 | 0 | 0 | 0 | 0 | 0 |

<sup>‡</sup>Indicates mRNA is circadian regulated in constant light according to analysis in Romanowski et al. (2020) The Plant Journal

### RVE8, LNK1, and COR27/28 form a protein complex

We used a yeast 2-hybrid system to validate the interactions identified in our APMS between RVE8/LNK1/LNK2 with COR27/28. Surprisingly, we did not see a positive interaction between these components when using a binary yeast 2-hybrid (**Fig. S4)**. Since APMS can identify both direct and indirect protein-protein interactions, we hypothesized that RVE8-LNK1/2-COR27/28 could be forming a protein complex where the CORs can only bind when both RVE8 and LNK1 are present. To test this, we used a yeast 3-hybrid system in which a linker protein is expressed in addition to the bait and prey proteins. We used N- and C-terminal truncations of LNK1 since full-length LNK1 autoactivates in yeast, as has been shown previously and here **(Fig. S5**) (Xie et al., 2014). Using this method, we found that yeast expressing RVE8, the C-terminus of LNK1, and COR27 or COR28 were able to grow on selective media in a higher order complex (**Fig. 3**). Yeast strains where COR27 or COR28 was paired with either LNK1 or RVE8 alone were unable to grow on selective media, indicating that indeed all three components must be present for the CORs to bind (**Fig. 3, S4**). We also confirmed that RVE8 interacts with the C-terminus of LNK1 (**Fig. S4**), in agreement with previous studies (Xie et al., 2014). In combination with our time-of-day APMS, these results show the CORs interact with RVE8/LNK1 in a complex that is present at ZT9.

**Figure 3.**
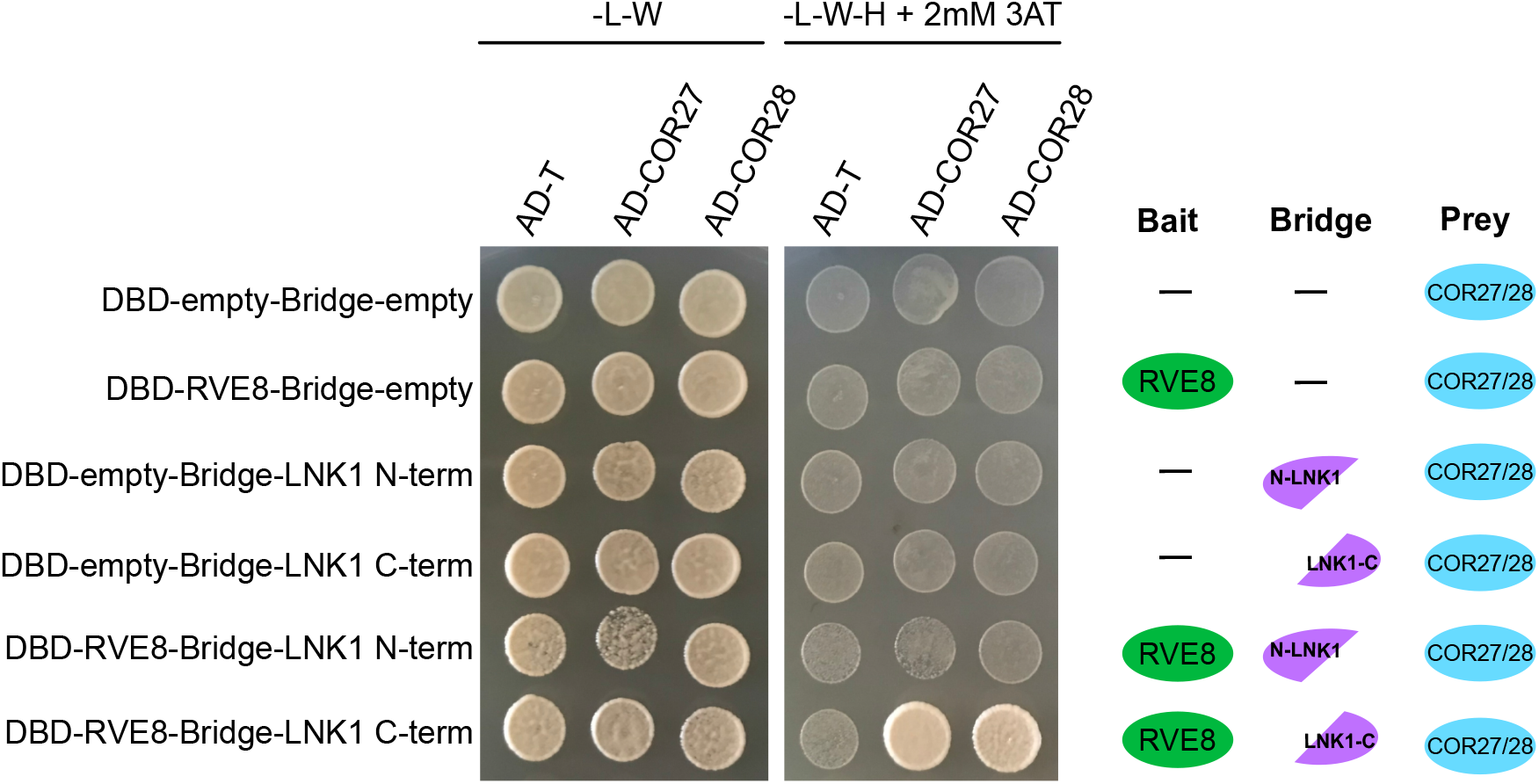
COR27/28 interact with RVE8/LNK1 in a yeast 3-hybrid system. Yeast strains Y2H Gold or Y187 expressing pBridge (GAL4-DBD and a Bridge protein) or pGADT7 (GAL4-AD), respectively, were mated and plated onto selective media. Successful matings can grow on - Leucine/-Tryptophan media (-L-W) while positive interactors can grow on -Leucine/-Tryptophan/- Histidine + 2mM 3-amino-1,2,4-triazole (3AT) (-L-W-H + 2mM 3AT). A graphical depiction of different combinations is shown to the right. AD-T (large T-antigen protein) is a negative control for prey interactions. Experiment was repeated at least twice.

### COR27/28 alter diurnal RVE8 protein abundance patterns and antagonize activation of the RVE8 target gene *TOC1*

We next sought to determine the biological relevance of the RVE8-LNK1/2-COR27/28 interaction. COR27/28 are post-translationally regulated via degradation by 26S proteasome (Kahle et al., 2020; Li et al., 2020; Zhu et al., 2020). As COR27/28 were identified as ZT9-specific RVE8-HFC interactors (**Table 2**), we hypothesized that COR27/28 target the RVE8-LNK complex for degradation in the evening, thus blocking expression of RVE8 target genes late in the day. To determine if RVE8-HFC abundance patterns are driven by a post-translational mechanism, we examined protein abundance of RVE8-HFC in seedlings treated with either the 26S proteasome inhibitor bortezomib (bortz) or DMSO (mock). The mock treated seedlings showed the typical pattern for RVE8-HFC protein abundance (**Fig. 1F**) with decreasing RVE8-HFC from ZT6 to ZT15 (**Fig. S6**). Treatment with bortz led to increased RVE8-HFC accumulation during this time frame, indicating 26S-proteasome degradation is involved in the observed decrease of RVE8-HFC from ZT6 to ZT15 (**Fig. S6**).

Next, we tested if COR27 and COR28 regulate RVE8 protein abundance by examining cyclic protein abundance in RVE8p::RVE8-HFC versus RVE8p::RVE8-HFC in *cor27-2 cor28-2*. While RVE8-HFC abundance in the wild-type background exhibits rhythmic protein abundance with peak protein levels at ZT6, RVE8-HFC abundance is significantly higher in the *cor27-2 cor28-2* background during the evening and nighttime hours (**Fig. 4A-C**). This result is consistent with the hypothesis that in the absence of COR27/28, RVE8-HFC should be stabilized specifically in the evening—when it would normally be degraded through its interaction with the CORs. As the circadian rhythm of *RVE8* mRNA expression under LD cycles was shown to be unchanged in the *cor27-2 cor28-2* background (Wang et al., 2017), our results indicate that COR27/28 regulate RVE8-HFC protein abundance at the post-translational level.

**Figure 4.**
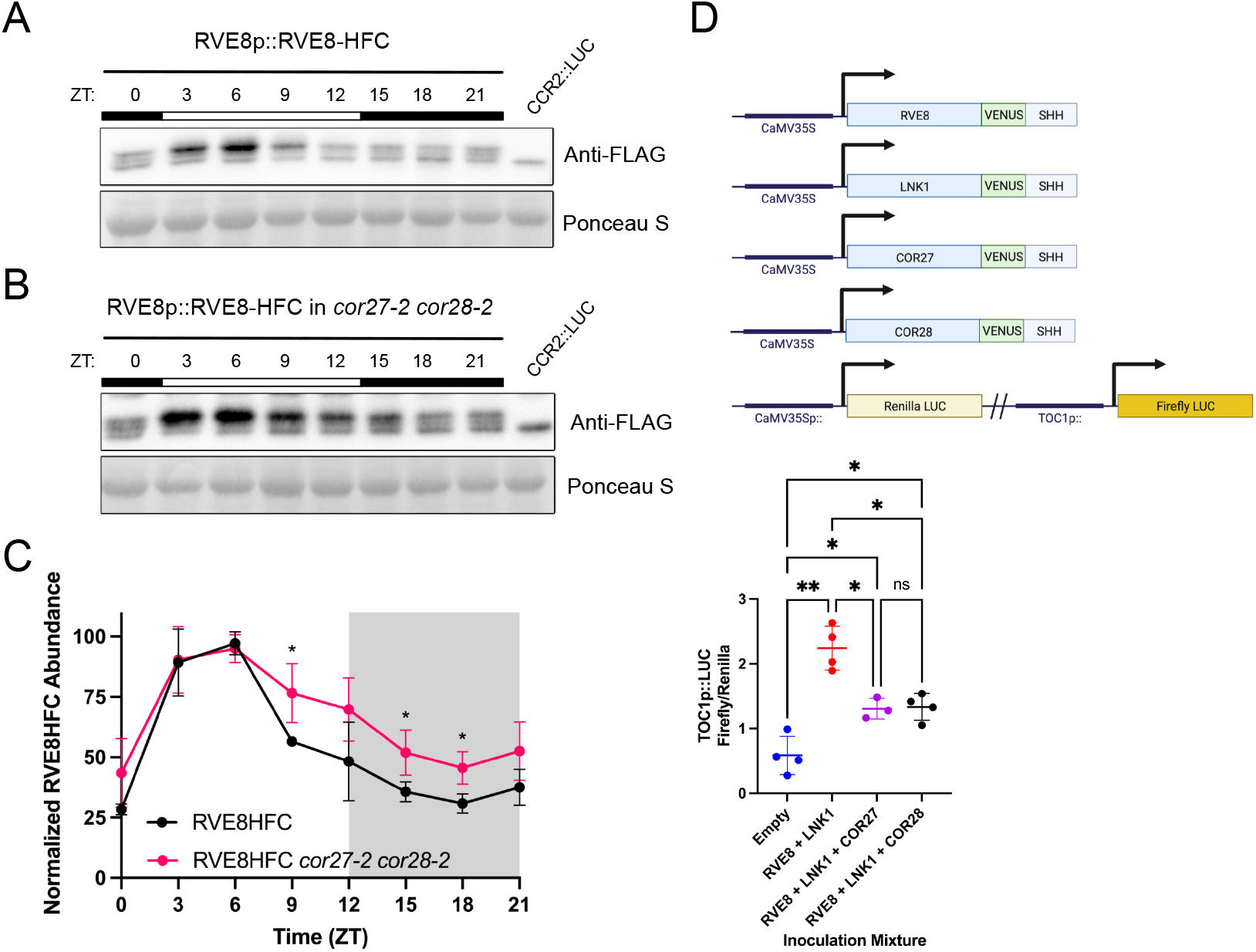
COR27/28 alter RVE8-HFC protein abundance patterns and inhibit RVE8/LNK1-mediated activation of *TOC1*. (A-B) 24-hour protein expression patterns of RVE8-HFC in wild type (A) or *cor27-2 cor28-2* (B) backgrounds analyzed by Western blot. Tissue was collected every 3 hours from 12-day-old plants grown under 12 hr light: 12 hr dark 22 °C conditions. Anti-FLAG antibody was used to detect RVE8-HFC and Ponceau S staining was used to show loading. White and black bars indicate lights-on and lights-off, respectively. Col-0 CCR2::LUC was used as the negative control. (C) Densitometry quantification of (A) and (B) RVE8-HFC 24-hour abundance normalized to Ponceau S in wild type and *cor27-2 cor28-2* backgrounds. Points represent the average normalized RVE8-HFC abundance from 3 (WT) or 4 (*cor27-2 cor28-2*) independent bioreps. Asterisks indicate significant differences between genotypes based on Welch’s t-test (* p< 0.05). Error bars = SD. (D) Dual luciferase assay in 3–4-week-old *Nicotiana benthamiana*. Schematic of expression constructs infiltrated are shown at the top (SHH = 2X-StrepII-HA-His**6** tag). Luminescence from a dual firefly/renilla luciferase reporter was measured after coinfection with 35S::RVE8-VENUS-SHH, 35S::LNK1-VENUS-SHH, 35S::COR27-VENUS-SHH, or 35S::COR28-VENUS-SHH. Luminescence was normalized to constitutively expressed renilla luciferase luminescence to control for infection efficiency. Points represent the normalized luminescence from 3-4 independent experiments with N=12. Mean normalized luminescence is indicated by the crosshair symbol and error bars = SD. Asterisks indicate significant differences by unpaired t-test with Welch correction (ns= not significant, * p< 0.05, ** p< 0.01). ZT= Zeitgeber Time. Empty = reporter alone.

We then tested the effect of the CORs on RVE8/LNK1 transcriptional activity using a transactivation assay in *Nicotiana benthamiana* (**Fig. 4D**). RVE8 binds to the evening element cis-regulatory motif in the *TOC1* promoter to activate its expression (Rawat et al., 2011). When LNK1 and RVE8 were transiently expressed together in *N. benthamiana* along with a *TOC1* promoter-driven luciferase reporter, we observed activation of the reporter, as expected (**Fig. 4D**). When COR27 or COR28 was added to the inoculation cocktail, activation of the reporter was reduced, indicating that the CORs antagonize RVE8/LNK1 transcriptional activity *in vivo* (**Fig 4D**). Taken together, our results indicate that the RVE8-COR27/28-LNK1/2 interaction serves to block activation of RVE8 target genes via degradation of RVE8 in the evening.

### The RVEs are important for cold temperature induction of COR27/28

*COR27/28* contain evening elements in their promoters that are important for their cold induction and could be targets of RVE8 transcriptional regulation (Mikkelsen and Thomashow, 2009; Wang et al., 2017). Additionally, *COR27/28* are significantly upregulated in an inducible RVE8:GR line according to a previously published RNA-seq dataset (Hsu et al., 2013b). Both COR27/28 and RVE4/8 regulate cold tolerance in Arabidopsis; *COR27/28* expression is induced by cold temperature (16 *°C* and 4 °C) within 3 hours and the *cor27-1 cor28-2* loss-of-function mutant shows increased freezing tolerance, suggesting these genes are negative regulators of the plant’s response to freezing temperatures (Li et al., 2016). In contrast, RVE4/8 are activators of cold tolerance (Kidokoro et al., 2021). Upon cold treatment (4°C for 3 hours), RVE4/8 localize to the nucleus and upregulate *DREB1A* to promote freezing tolerance (Kidokoro et al., 2021).

To determine if the RVE transcription factors are regulators of *COR27/28* cold-induction, we examined *COR27/28* expression at 22 °C and 4 °C in Col-0, *rve8-1, rve34568*, and *lnkQ* mutants. We found that *COR27/28* cold-induction was greatly attenuated in *rve34568* and *lnkQ* mutants, consistent with the *CORs* being targets of the RVE-LNK transcriptional complex (**Fig. S7A-B)**. The absence of an effect in the *rve8-1* single mutant suggests there is redundancy among the *RVE* family in the regulation of *COR27*/*28*. Indeed, we found that the LNKs coprecipitated RVE3/4/5/6/8 in our APMS (**Tables 1 and 2**), suggesting multiple RVE/LNK complexes could influence the regulation of the *CORs*. Interestingly, we saw little effect of RVEs/LNKs on *COR27/28* expression at 22 °C at ZT12 (**Fig. S7C-D**), suggesting these clock factors only have an effect under cold stress or that there may be a greater effect on expression at 22 °C at a different time of day.

### LNK1 and LNK2 are important for temperature entrainment of the clock

The enrichment of temperature response GO terms among the list of coprecipitated proteins in our APMS (**Fig. 2A**), as well as the existing evidence linking RVE8 to temperature regulation (Blair et al., 2019; Kidokoro et al., 2021) prompted us to investigate whether *LNK1/2* are important for temperature input to the clock. While light is the primary entrainment cue for the plant clock, daily temperature cycles are known to be another major environmental input cue (Devlin and Kay, 2001; Salomé and Robertson Mcclung, 2005; Avello et al., 2019). To examine temperature entrainment, we examined rhythms from a CCA1::LUC reporter in wild type and *lnk1-1, lnk2-4*, and *lnk1-1 lnk2-4* mutant plants that were first grown under constant light and then transferred into a temperature entrainment condition. Under constant light, the *lnk* mutants exhibited their canonical long period mutant phenotype (Rugnone et al., 2013) (**Fig. 5**). Upon entering a temperature entrainment condition of 12 hr 20 °C: 12 hr 22 °C, the *lnk1/2* mutants were unable to resynchronize their circadian rhythms to that of wild type (**Fig. 5A-B**). This defect was ameliorated when the difference between the minimum and maximum temperature was increased from 2 °C to 4 °C; when provided temperature cycles of 12 hr 18 °C: 12 hr 22 °C, most *lnk* mutants were able to realign with the wild-type acrophase (peak reporter expression) by the third day of temperature entrainment (**Fig. 5 C-D**). However, this resynchronization was still slower than when the *lnk* mutants were provided with photocycles—upon the transition from constant light to LD cycles, all mutants were able to immediately re-align their rhythms to wild type, indicating that the *lnk* mutants are specifically impaired in their ability to use temperature as an entrainment cue (**Fig. 5E-F**).

**Figure 5.**
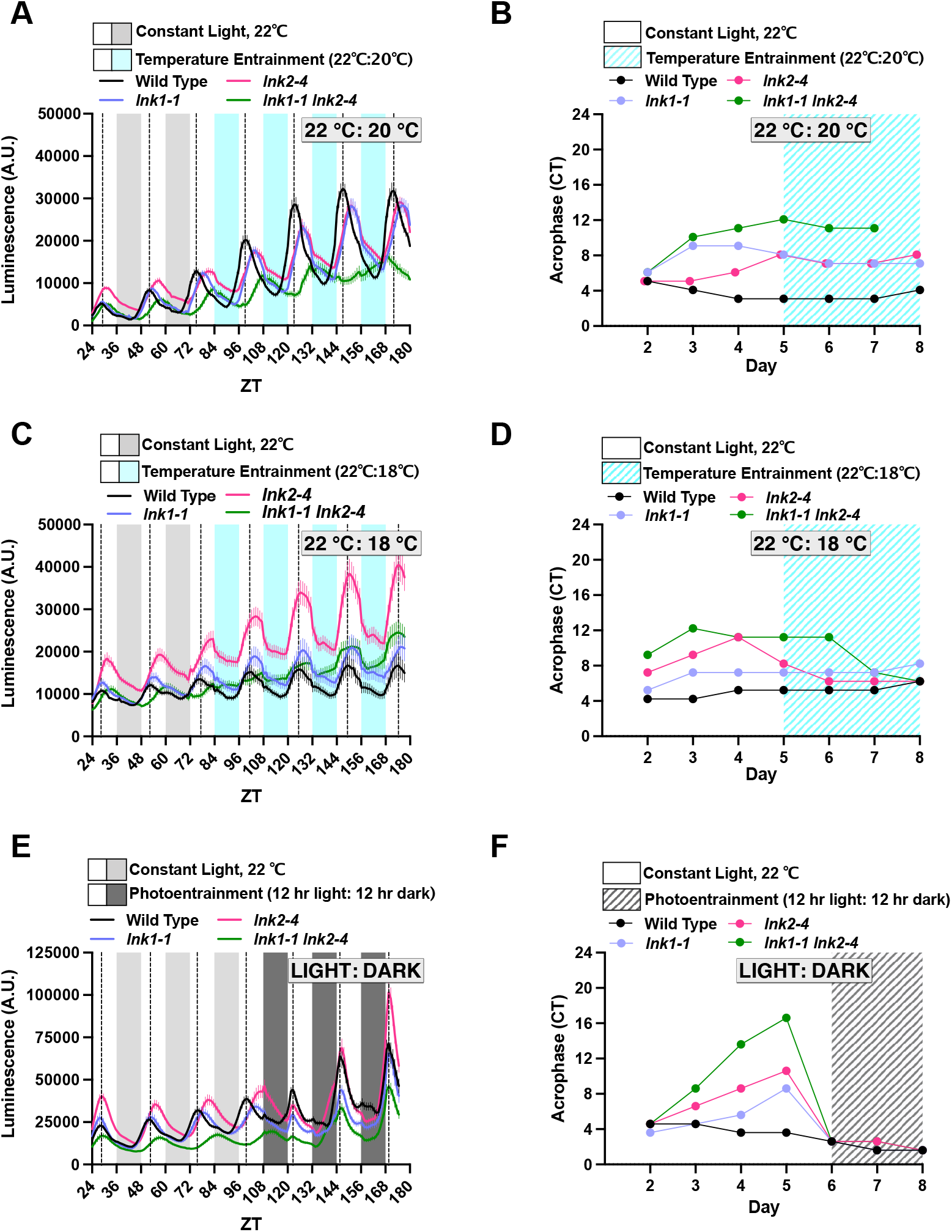
LNK1/2 are important for temperature entrainment of the clock. (A,C,E) Plants were grown for 7 days under 12 hr light: 12 hr dark 22 °C conditions for initial entrainment. On day 7, seedlings were transferred to imaging chamber and luminescence was measured for at least 3 days in continuous light and temperature (22 °C) before the chamber was switched to either a temperature- (A,C) or photo- (E) entrainment program. Temperature entrainment consisted of a day temperature of 22 °C and nighttime temperature of 20 °C (A) or 18 °C (C). Photoentrainment consisted of 12 hr light followed by 12 hr darkness (22 °C). Lines represent the average luminescence from n=16 seedlings with errors bars = SEM. Vertical dotted lines correspond to the acrophase, or time of peak reporter expression, of the CCA1::LUC reporter in wild type plants. (B,D,F) Acrophase is plotted for each genotype for each day of imaging in constant light and the temperature entrainment condition (B, D) or under photoentrainment (F). Each point represents the acrophase of the averaged luminescence trace shown in (A,C,E). CT = Circadian Time. A.U. = Arbitrary Units. ZT=Zeitgeber Time.

The temperature entrainment programs used in **Figure 5A-D** are non-ramping, meaning the temperature shifts immediately from the cool to warm temperatures. To better simulate environmental conditions, we also employed a ramping, natural temperature entrainment which gradually oscillates between a low temperature of 16 *°C* and a high of 22 °C. We observed a similar delay in the ability of the *lnk* mutants to assimilate to wild-type acrophase under natural temperature cycles, demonstrating that this defect is not a byproduct of non-ramping temperature changes (**Fig. S8**).

As the LNKs form a four-member family, we also examined whether LNK3/4 play a role in temperature entrainment. The *lnk3-1 lnk4-1* double mutant showed little difference from wild-type rhythms under constant light nor temperature entrainment, indicating LNK1/2 are the primary family members important for temperature entrainment (**Fig. S9**). In summary, we have demonstrated a previously unknown role for LNK1/2 in temperature entrainment of the clock.

## Discussion

Daily and seasonal temperature cycles are important cues for the entrainment of the plant circadian clock (Salomé and Clung, 2005). In parallel to this, the clock is essential for proper response to temperature stimuli (Salomé and Robertson Mcclung, 2005; Thines and Harmon, 2010). In this study, we have identified a novel, time-of-day-specific interaction between two established components of the circadian and temperature response pathways: the circadian clock transcriptional activation complex containing RVE8 and LNK1/LNK2 and the cold response proteins COR27/COR28. Previous studies have demonstrated that RVE8 and COR27/COR28 both regulate the transcription of the master cold response regulator *DREB1A* and the core circadian oscillator genes *PRR5* and *TOC1;* however, RVE8 acts as a transcriptional activator of these targets while the CORs act as repressors (Rawat et al., 2011; Li et al., 2016; Wang et al., 2017; Kidokoro et al., 2021). In addition to sharing transcriptional targets, RVE8 and COR27/COR28 also affect similar phenotypes, including period lengthening in the null or knock-down mutants and regulation of photoperiodic flowering time (Rawat et al., 2011; Li et al., 2016). Despite these established overlaps in function between the RVE8-LNK1/LNK2 complex and the CORs, a mechanistic connection between these factors has until now been lacking. In this study, we have demonstrated that COR27/COR28 physically interact with and regulate the protein stability of the RVE8-LNK1/LNK2 complex in the evening and that the CORs antagonize RVE8/LNK1-mediated activation of *TOC1* expression.

Our time-of-day-specific APMS experiments demonstrated that RVE8, LNK1, and LNK2 interact with different protein partners at ZT5 versus four hours later at ZT9 (**Fig. 2A-D**). LNK1 and RVE8 interacted with more protein partners at the later timepoint, ZT9, while LNK2 coprecipitated more interactors at ZT5 (**Fig. 2B-D)**. For LNK1 and LNK2, their time of peak protein abundance (**Fig. 1D**) aligned with the time of day when they coprecipitated the most interactors (**Fig. 2B-C**), suggesting that increased abundance of these clock bait proteins led to an increased number of captured interactions. Interestingly, while our 24-hour time course Western blots showed a higher abundance of RVE8-HFC at ZT5, we coprecipitated more interactors at ZT9 than at ZT5. This might indicate that even though protein levels of RVE8-HFC are lower at ZT9, perhaps there is an important bridge protein expressed in the evening that links in RVE8-HFC interactors only in the evening. Alternatively, perhaps there are more RVE8-HFC protein interacting partners expressed at ZT9 than at ZT5. By performing APMS at two different time points, we have established that these circadian clock proteins interact with different partners depending on the time of day.

For example, COR27, COR28, COP1, and SPA1 were coprecipitated with RVE8/LNK1/LNK2 at ZT9 but not ZT5 (**Tables 1-2**). We have considered the following hypotheses for what is driving this time-of-day-specific interaction: 1) The diurnal expression patterns of these components produces high gene expression overlap at ZT9 but not ZT5, 2) There is a third protein component that is expressed at ZT9 that allows for the interaction between these factors via bridging or by inducing a conformational change in one of the participating proteins, or 3) APMS is not an exclusionary method and could simply have not detected a low abundance peptide that was coprecipitated at ZT5. When we examined the LD mRNA expression patterns for *COR27, COR28, COP1*, and *SPA1*, we found that COR27 and COR28 are most likely ZT9-specific interactors due to their mRNA expression levels having a higher overlap with RVE8/LNK1/LNK2-HFC protein abundance at ZT9 (**Fig. S3**). Indeed, the *CORs* have very low mRNA expression at ZT5 and thus are likely absent from the cell and not interacting with the RVE8-LNK1/2 proteins (**Fig. S3**). COP1 and SPA1, in contrast, do not show higher expression overlap with RVE8-HFC at ZT9 over ZT5 (**Fig. S3**). We instead think it is possible that COP1/SPA1 could be recruited to RVE8 via COR27/COR28 and thus can only be coprecipitated at ZT9 (hypothesis #2). However, future studies are needed to validate this possibility.

As COR27/28 are post-translationally regulated by 26S proteasome-mediated degradation (Kahle et al., 2020; Li et al., 2020; Zhu et al., 2020), we predicted that the interaction between RVE8/LNK1/LNK2 and COR27/28 could function to target the circadian transcriptional module for degradation in the evening. We found that RVE8-HFC cyclic protein abundance patterns were disrupted in a *cor27-2 cor28-2* mutant background, with higher RVE8-HFC levels observed specifically during the evening and nighttime hours (**Fig. 4A-C**). This suggests that COR27/28 are important for degradation of RVE8 in the evening. As COP1/SPA1 were also identified as ZT9-specific RVE8 binding proteins, we suggest that the CORs recruit the COP1-SPA1 E3 ubiquitin ligase complex to RVE8-LNK1/2 to target it for degradation by the proteasome, though this has yet to be directly tested. We also coprecipitated *UBIQUITIN-SPECIFIC PROTEASE 12* (*UBP12*) and *UBP13* and the E3 ubiquitin ligases *PLANT U-BOX 12* (*PUB12*) and *PUB13* in RVE8/LNK1/LNK2 APMS experiments and these factors may also play a role in time-of-day-specific complex degradation (**Tables 1-2, Dataset S1**) (Zhou et al., 2021). In tobacco transactivation assays, we observed that presence of COR27/28 reduced the ability of RVE8-LNK1 to activate the expression of a *TOC1* promoter-driven reporter, demonstrating that the CORs have an antagonistic effect on the transcriptional activity of this circadian module (**Fig. 4D**).

The CORs do not have identifiable DNA-binding domains and do not bind to DNA *in vitro* (Li et al., 2020); therefore, the CORs must work with a DNA-binding protein to affect transcription of their target genes. Previous work supported this hypothesis by showing that COR27/28 interact with the major photomorphogenic transcription factor ELONGATED HYPOCOTYL 5 (HY5) and regulate some of the same HY5 target loci (Li et al., 2020). Perhaps a similar mechanism is at work here, with the CORs interacting with the RVE-LNK complex to alter its transcriptional activity. The mechanism behind how the CORs change or potentially change the activity of these transcription factors is an open question.

Finally, as *COR27/28* expression is induced under cold stress and RVE8 accumulates in the nucleus upon cold treatment, this presents an interesting possibility that the interaction between RVE8 and the CORs could serve to connect cold temperature response and the circadian clock. Notably, COR27/28 and RVE8 oppositely regulate freezing tolerance; the CORs repress expression of *DREB1A* to decrease freezing tolerance while RVE4/8 activate *DREB1A* expression (Li et al., 2016; Kidokoro et al., 2021). Thus, we anticipate that the interaction between the CORs and the RVE8-LNK complex is antagonistic in its nature.

In summary, we used affinity purification-mass spectrometry (APMS) to identify novel circadian-associated proteins using the RVE8/LNK1/LNK2 core circadian oscillator proteins as baits. By performing APMS at two time points during the 24-hour cycle, we identified time-of-day-specific interactors, including COR27 and COR28, which only coprecipitated with these three clock baits at the later timepoint, ZT9 **(Fig. 6A, Tables 1 and 2**). The obligate higher order nature of this complex that we established using a yeast 3-hybrid demonstrates a powerful advantage of using an *in vivo* method like APMS over another screening system—screens such as the yeast 2-hybrid library system can only identify binary interactions and thus would never have identified the interaction described here between RVE8, the C-terminus of LNK1, and COR27/28. Taken together, we propose the following model **(Fig. 6B**): In the morning–early afternoon, when the *CORs* are not expressed, the RVE8-LNK1/2 complex is free to perform its canonical duty as an activating force in the circadian oscillator and in cold tolerance. As evening approaches, *COR27/28* expression rises and the RVE8-LNK1/2-COR27/28 complex is formed, which antagonizes RVE8-LNK1/2 transcriptional activity via regulating RVE8 protein abundance. Future studies examining this complex’s role in circadian and cold tolerance phenotypes will be of great interest.

**Figure 6.**
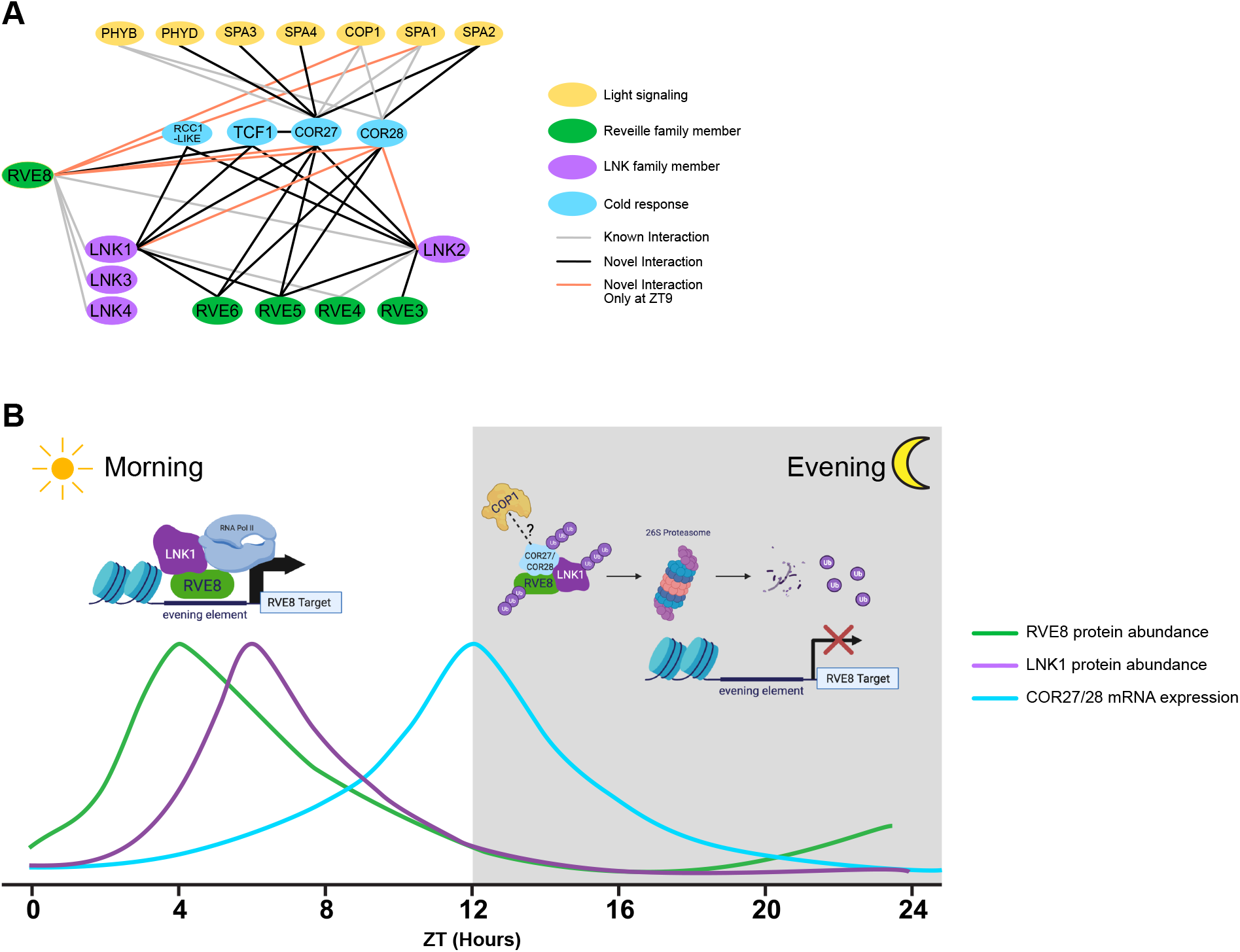
The RVE8-LNK1/2-COR27/28 complex is a novel post-translational regulatory mechanism in the circadian clock. (A) Protein interaction network compiled from APMS experiments using RVE8-HFC, LNK1-HFC, LNK2-HFC, YFP-COR27, and GFP-COR28 as bait proteins at ZT5 and ZT9. Black lines indicate novel interactions identified in this study, grey lines show previously published interactions validated in this study, and orange lines show novel interactions that were identified only at ZT9. (B) Model of hypothesized role of the RVE-LNK-COR interaction during a 24-hour period. In the morning, RVE8-LNK1/2 interact to coactivate the expression of target genes such as evening-phased circadian clock genes and cold-response genes. Towards the evening, COR27/28 are expressed and interact with the RVE8-LNK1/2 complex, potentially recruiting a ubiquitin E3 ligase such as COP1 to target the entire complex for degradation by the 26S proteasome, thus blocking activation of RVE8 targets in the evening. Green and purple lines show approximate protein abundance patterns of RVE8 and LNK1, respectively, while the blue line shows approximate *COR27/28* mRNA expression.

## Methods

### Plant Materials

T-DNA disrupted lines used in this study: rve8-l (SALK_053482C), lnk1-1 (SALK_024353), lnk2-1 (GK_484F07), lnk2-4 (GK_484F07), lnk3-1 (SALK_085551C), lnk4-1 (GK_846C06), cor27-2 (SALK_042072C), and cor28-2 (SALK_137155C) (Alonso et al., 2003). The lnkQ CCA1::LUC line was generated by transforming the lnkQ mutant background (de Leone et al., 2020) with a binary vector containing CCA1::LUC and Basta resistance (from Harmer Lab). The *lnk3-1 lnk4-1* CCA1::LUC line was generated by crossing *lnk3-1 lnk4-1* to the CCA1::Luc reporter. The 35S::YFP-COR27 and 35S::GFP-COR28 lines were described previously (Li et al., 2016) and generously shared with us by Dr. Hongtao Liu. The rve8-1 CCR2::LUC line was described previously (Rawat et al., 2011) and generously shared with us by Dr. Stacey Harmer. The lnk1-1 CCA1::LUC, lnk2-4 CCA1::LUC, and lnk1-1 lnk2-4 CCA1::LUC lines were a generous gift from Dr. Xiaodong Xu (Xie et al., 2014). All plants used were in the Col-0 background.

Seeds were gas sterilized and plated on 1/2X Murashige and Skoog basal salt medium with 0.8% agar + 1% (w/v) sucrose. After stratification for 2 days, plates were transferred to a Percival incubator (Percival-Scientific, Perry, IA) set to a constant temperature of 22 °C. Light entrainment was 12 hr light/12 hr dark (LD) cycles, with light supplied at 80 μmol/m^2^/s. 24-hour tissue collections were performed under white light during the daytime timepoints and under dim green light during the nighttime timepoints.

### Generation of Epitope-tagged Lines and Plasmid Construction

To generate pB7-RVE8p::RVE8-HFC, RVE8 was cloned from genomic DNA without the stop codon using primers pDAN1127 and pDAN1128 (**Table S2**) and cloned into NotI/AscI-digested pENTR-MCS through In-Fusion HD cloning (Clontech, Mountain View, California). pENTR-RVE8-no stop was then recombined using LR Clonase (Thermo Fisher Scientific, Waltham, Massachusetts) into pB7-HFC (Huang et al., 2016a), which contains the 6X-HIS 3X-FLAG C-terminal tag, to generate pB7-RVE8-HFC. To generate the endogenous promoter driven line, the sequence upstream of the RVE8 transcription start site to the stop codon of the upstream gene was cloned (945 bases) using primers pDAN1129 and pDAN1130 (**Table S2**). The 35S Cauliflower Mosaic Virus (CaMV35S) promoter was excised from pB7-RVE8-HFC via PmeI/SpeI digest and replaced with the RVE8 promoter fragment through In-Fusion HD cloning (Clontech, Mountain View, California) to generate pB7-RVE8P::RVE8-HFC. pB7 RVE8p::RVE8-HFC binary vector was transformed into rve8-1 CCR2::LUC (Rawat et al., 2011) by agrobacterium mediated transformation and positive transformants were identified through basta resistance (Clough and Bent, 1998).

To generate pH7WG2-LNK1p::LNK1-HFC and pH7WG2-LNK2p::LNK2-HFC, LNK1 and LNK2 coding sequences were cloned from cDNA without the stop codon using primers pDAN0990/pDAN0991 (LNK1) and pDAN1066/pDAN1067 (LNK2) (**Table S2**) and recombined into pENTR-MCS through dTOPO cloning or In-Fusion HD cloning (Contech, Mountain View, California), respectively. pENTR-LNK1-no stop and pENTR-LNK2-no stop were then recombined using LR Clonase (Thermo Fisher Scientific, Waltham, Massachusetts) into pB7-HFC to generate pB7-LNK1-HFC and pB7-LNK2-HFC. To make the endogenous promoter driven construct, the LNK1 promoter was cloned from the LNK1 transcription start site to the upstream gene’s 5’ UTR (1709 bp) using primers pDAN1016 and pDAN1017 (**Table S2**). This promoter fragment was swapped with CaMV35S via PmeI/SpeI digest and In-Fusion HD cloning (Clontech, Mountain View, California) to generate pB7-LNK1p::LNK1-HFC. Similarly, the LNK2 promoter was cloned from just before the start of the upstream gene through 142 bases into exon 4 from genomic DNA using primers pDAN1018 and pDAN1019 (**Table S2**) and inserted into pB7-HFC PmeI/BglII digest and In-Fusion HD cloning (Clontech, Mountain View, California) to generate pB7-LNK2p::LNK2-HFC. To make pH7WG2-LNK1p::LNK1-HFC and pH7WG2-LNK2p::LNK2-HFC, pB7-LNK1p::LNK1-HFC, pB7-LNK2p::LNK2-HFC, and pH7WG2 (Karimi et al., 2002) were digested with KpnI and AgeI and the resulting fragments were ligated. pH7-LNK1p::LNK1-HFC and pH7-LNK2p::LNK2-HFC binary vector were transformed into lnkQ CCA1::LUC by agrobacterium mediated transformation and positive transformants were identified through hygromycin resistance (Clough and Bent, 1998).

To make LNK1 truncations, the N-terminus of LNK1 from the start codon through amino acid 296 was cloned using primers pDAN1954/pDAN2010 (**Table S2**), adding a stop codon. The LNK1 C-terminal fragment was cloned using primers pDAN2011/pDAN1955 (**Table S2**) with the first amino acid starting at amino acid number 297. Gene fragments were recombined into pENTR-MCS through In-Fusion HD cloning (Clontech, Mountain View, California) to make pENTR-LNK1-N-term-STOP and pENTR-LNK1-C-term-STOP.

To generate pK7-VENUS (VEN)-2x-StrepII-HA-6X-His-C-terminus (SHHc), we first made pK7-SHHc by PCR amplifying the 2X-SII-HA-6X-His C-terminal (SHHc) tag from pB7-SHHc (Huang et al., 2016b) and digesting pK7FWG2 (Karimi et al., 2002) with BstXI and KpnI. The PCR fragment containing the SHHc tag was combined with the digested backbone using In-Fusion HD cloning (Clontech, Mountain View, California) to make pK7-SHHc. Venus was cloned from plasmid mVENUS C1 (Koushik et al., 2006) using primers pDAN0869 and pDAN0870 and recombined with pK7SHHc digested with AvrII using In-Fusion HD cloning (Clontech, Mountain View, California) to generate pK7-VEN-SHHc.

pENTR-no stop clones of COR27 and COR28 were generated by amplifying the coding sequences of COR27 (AT5G24900.1) and COR28 (AT4G33980.1) using primers pDAN1906/pDAN1908, and pDAN1909/pDAN1911, respectively (**Table S2**). The resulting amplicons were cloned into NotI/AscI-digested pENTR-MCS through In-Fusion HD cloning (Clontech, Mountain View, California) to make pENTR-COR27-no stop and pENTR-COR28-no stop. To generate pK7-RVE8-VEN-SHHc, pK7-LNK1-VEN-SHHc, pK7-COR27-VEN-SHHc, and pK7-COR28-VEN-SHHc, the pENTR-no stop versions of these genes were recombined to the pK7-VEN-SHHc binary vector using LR Clonase (Thermofisher). These C-terminally tagged proteins are driven from the CaMV35S promoter. To generate the dual luciferase reporter pGreenII 0800-LUC-TOC1p, 2098 bp of the TOC1 promoter was cloned using primers pDAN2735/pDAN2736 (**Table S2**) and inserted via In-Fusion HD cloning (Clontech, Mountain View, California) into the pGreenII 0800-LUC plasmid (Hellens et al., 2005) digested with BamHI. The resulting vector (pGreenII 0800-LUC-TOC1p) constitutively expresses renilla luciferase from the CaMV35S promoter and contains the gene for firefly luciferase driven by the TOC1 promoter.

To generate yeast 2-/3-hybrid vectors, the gene of interest was cloned from its pENTR-STOP template using primers pDAN2349/pDAN2350 (**Table S2**) and recombined into pGADT7 digested with EcoRI using In-Fusion HD cloning (Clontech, Mountain View, California). For cloning into pGBKT7, primers pDAN2347/pDAN2348 (**Table S2**) were used to clone off the pENTR-STOP template and recombine into BamHI-digested pGBKT7 using In-Fusion HD cloning (Clontech, Mountain View, California). For cloning into the pBridge vector (Clontech, Mountain View, California), the gene of interest was cloned from its pENTR-STOP template using primers pDAN2441/pDAN2442 (**Table S2**) and recombined into the first MCS of pBridge digested with EcoRI using In-Fusion HD cloning (Clontech, Mountain View, California) or using primers pDAN2443/pDAN2444 (**Table S2**) to recombine into the second MCS of pBridge digested with BglII using In-Fusion HD cloning (Clontech, Mountain View, California).

### Affinity Purification

Affinity purification was performed as detailed in Sorkin and Nusinow (2022). Briefly, affinity-tagged lines were plated on 1/2x MS + 1% Sucrose and grown for 10 days under LD 22 °C conditions. On day 10 of growth, tissue was harvested at either ZT5 or ZT9. To extract protein, powdered tissue was resuspended in SII buffer (100 mM sodium phosphate, pH 8.0, 150 mM NaCl, 5 mM EDTA, 5 mM EGTA, 0.1% Triton X-100, 1 mM PMSF, 1x protease inhibitor mixture (Roche, Basel, Switzerland), 1x Phosphatase Inhibitors II & III (Sigma-Aldrich), and 5 μM MG132 (Peptides International, Louisville, KY)) and sonicated using a duty cycle of 20 s (2 s on, 2 s off, total of 40 s) at 50% power. Extracts were clarified of cellular debris through 2x centrifugation for 10 min at ≥20,000 × g at 4 °C.

For HFC-tagged samples, clarified extracts were incubated with FLAG-M2-conjugated Protein G Dynabeads (Thermo Fisher Scientific, Waltham, Massachusetts) for one hour. Captured proteins were eluted off FLAG beads using 500 μg/mL 3x-FLAG peptide (Sigma-Aldrich). Eluted proteins were then incubated with Dynabeads His-Tag Isolation and Pulldown (Thermo Fisher Scientific, Waltham, Massachusetts) for 20 minutes and then washed 5 x 1 minute in His-tag Isolation Buffer (100 mM Na-phosphate, pH 8.0, 150 mM NaCl, 0.025% Triton X-100). Washed bead pellet was washed 4x in 25mM ammonium bicarbonate and flash frozen in liquid N_2_.

For YFP-COR27 and GFP-COR28, clarified extracts were incubated with GFP-TRAP Magnetic Agarose affinity beads (ChromoTek GmbH, Planegg-Martinsried, Germany) for one hour. Captured proteins were washed 3 x 1 minute in His-tag Isolation Buffer (100 mM Na-phosphate, pH 8.0, 150 mM NaCl, 0.025% Triton X-100) and 4x in 25mM ammonium bicarbonate and then flash frozen in liquid N_2_.

### LC-MS/MS analysis of AP samples

Samples on affinity beads were resuspended in 50 mM ammonium bicarbonate, reduced (10 mM TCEP) and alkylated (25 mM Iodoacetamide) followed by digestion with Tryspin at 37°C overnight. Digest was separated from beads using a magnetic stand and acidified with 1%TFA before cleaned up with C18 tip (Thermo Fisher Scientific, Waltham, Massachusetts). The extracted peptides were dried down and each sample was resuspended in 10 μL 5% ACN/0.1% FA. 5 μL was analyzed by LC-MS with a Dionex RSLCnano HPLC coupled to a Orbitrap Fusion Lumos mass spectrometer (Thermo Fisher Scientific, Waltham, Massachusetts) using a 2h gradient. Peptides were resolved using 75 μm x 50 cm PepMap C18 column (Thermo Fisher Scientific, Waltham, Massachusetts).

Peptides were eluted at 300 nL/min from a 75 μm x 50 cm PepMap C18 column (Thermo Scientific) using the following gradient: Time = 0–4 min, 2% B isocratic; 4–8 min, 2-10% B; 8–83 min, 10–25% B; 83–97 min, 25-50% B; 97–105 min, 50-98%. Mobile phase consisted of A, 0.1% formic acid; mobile phase B, 0.1% formic acid in acetonitrile. The instrument was operated in the data-dependent acquisition mode in which each MS1 scan was followed by Higher-energy collisional dissociation (HCD) of as many precursor ions in 2 second cycle (Top Speed method). The mass range for the MS1 done using the FTMS was 365 to 1800 m/z with resolving power set to 60,000 @ 400 m/z and the automatic gain control (AGC) target set to 1,000,000 ions with a maximum fill time of 100 ms. The selected precursors were fragmented in the ion trap using an isolation window of 1.5 m/z, an AGC target value of 10,000 ions, a maximum fill time of 100 ms, a normalized collision energy of 35 and activation time of 30 ms. Dynamic exclusion was performed with a repeat count of 1, exclusion duration of 30 s, and a minimum MS ion count for triggering MS/MS set to 5000 counts.

### AP-MS Data Analysis

MS data were converted into mgf. Database searches were done using Mascot (Matrix Science, London, UK; v.2.5.0) using the TAIR10 database (20101214, 35,386 entries) and the cRAP database (http://www.thegpm.org/cRAP/) and assuming the digestion enzyme trypsin and 2 missed cleavages. Mascot was searched with a fragment ion mass tolerance of 0.60 Da and a parent ion tolerance of 10 ppm. Oxidation of methionine and carbamidomethyl of cysteine were specified in Mascot as variable modifications. Scaffold (Proteome Software Inc., Portland, OR; v.4.8) was used to validate MS/MS based peptide and protein identifications. Peptide identifications were accepted if they could be established at greater than 95.0% probability by the Peptide Prophet algorithm (Keller et al., 2002) with Scaffold delta-mass correction. The Scaffold Local FDR was used and only peptides probabilities with FDR <1% were used for further analysis. Protein identifications were accepted if they could be established at greater than 99.9% probability as assigned by the Protein Prophet algorithm (Nesvizhskii et al., 2003). Proteins that contained similar peptides and could not be differentiated based on MS/MS analysis alone were grouped to satisfy the principles of parsimony. Proteins sharing significant peptide evidence were grouped into clusters. Only the proteins identified with ≥2 unique peptides were further used in the analysis, except when proteins with only one peptide were identified in more than one replicate.

### Yeast 2-Hybrid (Y2H) and Yeast 3-Hybrid Assays

We used the GAL4-based Matchmaker Gold Yeast 2-Hybrid System (Clontech, Mountain View, California) for all Y2H and Y3H assays. All transformations were performed as detailed in the Yeast Protocols Handbook (Clontech, Mountain View, California). For Y2H, bait proteins were cloned into the pGBKT7 vector which encodes the GAL4 DNA binding domain and then transformed into the Y2H Gold strain (Clontech, Mountain View, California) and plated on SD/- Trp to select for positive transformants. Prey proteins were cloned into the pGADT7 vector which encodes the GAL4 activation domain, transformed into the Y187 strain (Clontech, Mountain View, California), and plated on SD/-Leu to select for positive transformants. All matings were performed as detailed in the Yeast Protocols Handbook (Clontech, Mountain View, California) using the 96-well plate format. Mated diploids were selected for on SD/-Leu/-Trp media. Single colonies of mated bait + prey strains were resuspended in YPDA and plated on SD/-Leu-Trp or SD/-Leu-Trp-His plates.

For Y3H, bait and linker proteins were cloned into the appropriate position of the pBridge vector (Clontech, Mountain View, California), which encodes a GAL4 DNA binding domain and a linker protein, transformed into the Y2H Gold strain, and plated on SD/-Trp to select for positive transformants. pBridge strains were mated with pGADT7 prey strains and plated on SD/-Trp/-Leu to select for diploids. Single colonies of mated strains were resuspended in YPDA plated on SD/- Leu-Trp or SD/-Leu-Trp-His plates.

### Luciferase Reporter Assays

Individual 6-day-old seedlings expressing a CCA1::LUC reporter grown under LD cycles at 22°C were arrayed on 1/2x MS + 1% Sucrose plates and sprayed with 5mM luciferin (GoldBio, Olivette, MO) prepared in 0.01% (v/v) Triton X-100 (Millipore Sigma-Aldrich, St. Louis, MO). Plants were transferred to an imaging chamber set to the appropriate free-run or entrainment program and images were taken every 60 minutes with an exposure of 10 minutes after a 3-minute delay after lights-off to diminish signal from delayed fluorescence using a Pixis 1024 CCD camera (Princeton Instruments, Trenton, NJ). Images were processed to measure luminescence from each plant using the Metamorph imaging software (Molecular Devices, Sunnyvale, CA). Circadian period was calculated using fast Fourier transformed nonlinear least squares (FFT-NLLS) (Plautz et al., 1997) using the Biological Rhythms Analysis Software System 3.0 (BRASS) available at http://www.amillar.org.

### *N. benthamiana* Transient Transformation

Transient transformation of 3-4 week-old *N. benthamiana* plants was performed as in (Lasierra and Prat, 2018). Briefly, overnight saturated cultures of *Agrobacterium tumefaciens* strain GV3101 carrying pGreenII 0800-LUC-TOC1p, pK7-RVE8-VEN-SHHc, pK7-LNK1-VEN-SHHc, pK7-COR27-VEN-SHHc, pK7-COR28-VEN-SHHc, or 35S::P19-HA (Chapman et al., 2004) were pelleted and resuspended in 5 mL of resuspension buffer (10mM MgSO_4_, 10mM MES (pH 5.8), 150 μM Acetosyringone) for 2-3 hours. Cultures were diluted to OD_600_= 0.4 in resuspension buffer and inoculation mixtures were prepared by mixing the selected constructs together with the volume of 35S::P19-HA being varied to ensure that an equal amount of agrobacteria was added to each mixture relative to the reporter, regardless of the total number of effectors being introduced. Mixtures were inoculated into one quadrant of a mature leaf per one mixture. Four different mixtures could be inoculated into a single leaf. Three leaves per plant were inoculated and four plants were used for a total of 12 biological replicates per mixture.

### Dual-Luciferase Assay

The dual luciferase assay was performed using the Dual-Glo Luciferase Assay System (Promega, Madison, Wisconsin). Briefly, 3-4 week-old tobacco plants were inoculated with *Agrobacterium tumefaciens* expressing pGreenII 0800-LUC-TOC1p and a combination of other proteins: pK7-RVE8-VEN-2x-StrepII-HA-6X-His-C-terminus (SHHc), pK7-LNK1-VEN-SHHc, pK7-COR27-VEN-SHHc, or pK7-COR28-VEN-SHHc. This reporter firefly luciferase driven by the 3 leaf disks were collected per infiltration site from 3-day-post-infiltrated tobacco plants and frozen in liquid N_2_. Tissue was homogenized and resuspended in 200 μL of Cell Culture Lysis Reagent (100 mM potassium phosphate, pH 7.8, 1 mM EDTA, 7mM 2-mercaptoethanol, 1% Triton X-100, 10% glycerol). Lysates were centrifuged at max speed for 5 minutes and 5 μL of undiluted extract was used for the Dual Luciferase Assay input. 40 μL of Luciferase Assay Buffer was added to undiluted extract in a black 96-well plate and incubated for at least 10 minutes. Luminescence was measured over a 10-minute exposure using a Pixis 1024 CCD camera (Princeton Instruments, Trenton, NJ). 40 μL of Stop & Glo Reagent was added to wells to quench the firefly luciferase signal and provide the substrate for renilla luciferase. After at least 10 minutes incubation, luminescence was measured over a 10-minute exposure using the CCD camera. Firefly luciferase signal was divided by renilla signal to calculate normalized luminescence.

### Densitometry Analysis

Densitometry analysis was performed in FIJI (https://imagej.net/software/fiji/) on high resolution (600 dpi), greyscale images of Western blots captured with the same exposure time. Mean grey value was measured from ROIs of equal area for each protein band and for background regions as well as for loading controls (Ponceau S stain) and loading control background regions. Inverted pixel density of background regions was subtracted from the inverted pixel density of protein bands and loading controls to generate the net pixel density value. To calculate normalized abundance, the ratio of the net protein band value over the net loading control value was taken.

### Quantitative RT-PCR

Seedlings were gas sterilized and grown on 1/2x MS + 1% Sucrose plates with Whatman filter paper under 12 hr light: 12 hr dark, 22 °C conditions. On day 7 of growth at ZT10, plates were transferred to a different chamber set to either 22 °C or 4 °C for two hours. Tissue was collected at ZT12. Total RNA was extracted from powdered tissue using the RNeasy Plant Mini kit (Qiagen, Hilden, Germany). 1 μg of total RNA was used as the template to synthesize cDNA using the iScript RT-PCR kit (Bio-Rad, Carlsbad, CA). qPCR was performed with the SYBR Green I nucleic acid gel stain (Sigma-Aldrich) using a QuantStudio 5 Real-Time PCR System (ThermoFisher). PCR was set up as follows: 3 min at 95°C, followed by 40 cycles of 10 s at 95°C, 10 s at 55°C and 20 s at 72°C. A melting curve analysis was conducted right after all PCR cycles are done. APA1 (At1g11910), expression of which remain stable during the diurnal cycle, was used as the normalization control. Primers for qPCR are listed in **Table S2**.

## Acknowledgements

We acknowledge support by the National Science Foundation under Grant No. DBI-1827534 for acquisition of the Orbitrap Fusion Lumos LC-MS/MS and from the NIH award 5R01GM141374 to DAN. MLS was supported by NSF GRF award DGE-1745038 and by the William H. Danforth Plant Science Fellowship from the Donald Danforth Plant Science Center. This study was also supported by the German Research Foundation (DFG) under Germany’s Excellence Strategy (CIBSS - EXC 2189 – Project ID 390939984) and DFG grant HI 1369/6-1 to AH and by Agencia Nacional de Promoción Científica y Tecnológica (ANPCyT) to MJY.

## Author Contributions

MLS, NK, AH, MJY, AR, BSE and DAN designed the research project. The research was performed by MLS, ST, and RB. AR contributed new analysis to the paper. MLS, ST, and DAN wrote the paper.

---

**Supplemental Figure 1 mRNA expression patterns of *RVE8, LNK1*, or *LNK2* under photocycles (12 hr light: 12 hr dark).** White and dark grey shading indicates lights-on and lights-off, respectively. Microarray data from diurnal.mocklerlab.com.

**Supplemental Figure 2 Protein alignment of TCF1 (AT3G55580) and RCC1L (AT3G53830).** Protein sequences were aligned using the needle algorithm using the EBLOSUM62 matrix, a gap penalty of 10.0, and an extend penalty of 0.5. Sequences share 49.7% identity.

**Supplemental Figure 3 Comparison of HFC-tagged protein abundance with *COR27/28, COP1*, and *SPA1* mRNA expression profiles.** 24-hour (12 hr light: 12 hr dark, 22 °C (LDHH)) protein abundance (dark blue) is quantified from Western blots shown in Figure 1D-F. LDHH mRNA data from diurnal.mocklerlab.com (light blue) is overlayed. Vertical dotted lines show the time of day when tissue was collected for APMS. White and grey shading indicated lights-on and lights-off, respectively.

**Supplemental Figure 4 COR27/28 do not interact with RVE8 or LNK1 in a binary Y2H system** Yeast strains Y2H Gold or Y187 expressing pGBKT7 (Gal4-DBD) or pGADT7 (Gal4-AD), respectively, were mated and plated onto selective media. Successful matings were able to grow on -Leucine/-Tryptophan media (-L-W) while positive interactors can grow on -Leucine/- Tryptophan/-Histidine + 2mM 3-amino-1,2,4-triazole (3AT) (-L-W-H +3AT). Only the positive controls DBD-53 (p53) + AD-T (large T-antigen protein) and DBD-RVE8 + AD-LNK1 C-term show an interaction.

**Supplemental Figure 5 Full-length LNK1 auto-activates in yeast when paired with a DBD-containing protein** Yeast strains Y2H Gold or Y187 expressing pBridge (Gal4-DBD and a Bridge protein) or pGADT7 (Gal4-AD), respectively, were mated and plated onto selective media. Successful matings were able to grow on -Leucine/-Tryptophan media (-L-W). Full length LNK1 (bridge protein, no AD domain) paired with the transcription factor RVE8 (*) can aberrantly activate the expression of the histidine biosynthesis reporter, allowing it to grow on -Leucine/- Tryptophan/-Histidine (-L-W-H) when paired with the negative control large T-antigen protein (T). LNK1 N- and C-terminal truncations do not autoactivate.

**Supplemental Figure 6 RVE8-HFC protein abundance patterns are regulated by the 26S proteasome** (A) Representative Western blot showing protein expression patterns of RVE8-HFC plants treated with DMSO or 100 μM bortezomib. At ZT5, 12-day-old seedlings growing under 12 hr light: 12 hr dark, 22 °C conditions were immersed in 1/2X MS media containing either 100 μM bortezomib or DMSO. Tissue was collected every 3 hours starting at ZT6. RVE8-HFC was detected with anti-FLAG and Ponceau S staining was used to show loading. (B) Densitometry quantification of RVE8-HFC abundance in (A) normalized to Ponceau S. Points represent the average normalized RVE8-HFC abundance from 3 independent bioreps. Asterisks indicate significant differences between genotypes based on Welch’s t-test (* p<0.05). Error bars = SD. White and grey shading indicate lights-on and lights-off, respectively. ZT= Zeitgeber Time.

**Supplemental Figure 7 The RVEs and LNKs are important for cold induction of *COR27/28*.** Seedlings were grown on 1/2X MS + 1% sucrose for seven days under 12 hr light; 12 hr dark 22 °C conditions and then transferred at ZT10 to either 22 °C or 4 °C for two hours and tissue was collected at ZT12. (A-B) show the induction of *COR27/28* expression at 4 °C compared to 22 °C. Figures (C-D) show *COR27/28* expression levels at 22 °C. Expression was normalized to the endogenous control gene APA1. Bars show average expression with error bars = SD from 3 independent bioreps (points) for each genotype. Asterisks indicate significant differences as determined by Welch’s t-test (** p<0.01, *** P<0.001).

**Supplemental Figure 8 LNK1/2 mutants are also impaired in temperature entrainment under ramping temperature cycles** (A) Luminescence from 7-day-old plants entrained under 12 hr light: 12 hr dark, 22 °C conditions expressing a CCA1p::LUC reporter was imaged for at least 3 days in continuous light and temperature (22 °C) before the chamber was switched to a ramping temperature entrainment program that gradually oscillated between a low temperature of 16 °C at ZT16 and a high of 22 °C at ZT4. Lines represent the average luminescence from n=16 seedlings with errors bars = SEM. Vertical dotted lines correspond to the peak expression time (acrophase) of the CCA1p::LUC reporter in wild type plants. (B) Acrophase, or time of peak reporter expression, is plotted for each genotype for each day of imaging in constant light and the temperature entrainment condition. Each point represents the acrophase of the averaged luminescence trace shown in (A). CT = Circadian Time. A.U. = Arbitrary Units. ZT= Zeitgeber Time.

**Supplemental Figure 9 *lnk3/4* mutants are not impaired in temperature entrainment** (A) Luminescence from 7-day-old plants expressing a CCA1p::LUC reporter were grown for at least 3 days in continuous light and temperature (22 °C) before the chamber was switched to a ramping temperature entrainment program that gradually oscillates between a low temperature of 16 °C at ZT16 and a high of 22 °C at ZT4. Lines represent the average luminescence from n=16 seedlings with errors bars = SEM. Vertical dotted lines correspond to the peak expression time of the CCA1p::LUC reporter in wild type plants. (B) Acrophase, or time of peak reporter expression, is plotted for each genotype for each day of imaging in constant light and the temperature entrainment condition. Each point represents the acrophase of the averaged luminescence trace shown on the right. CT = Circadian Time. A.U. = Arbitrary Units. ZT=Zeitgeber Time.

**Supplemental Table 1 *AT3G53830* (*RCC1L*) is downregulated at 4 °C**. Data taken from Kidokoro et al. (2021) PNAS. Wild-type (Col-0) plants were transferred to 4 °C at LL2 (T=0; 2 hours after dawn) and tissue for RNA sequencing was collected at 3 hours and 12 hours after transfer to cold conditions. *RCC1L* is significantly downregulated after 12 hours under 4 °C treatment.

**Supplemental Table 2 Oligonucleotides used in this study.**

