## Supplemental Figure 1 for "COR27/28 Regulate the Evening Transcriptional Activity of the RVE8-LNK1/2 Circadian Complex"

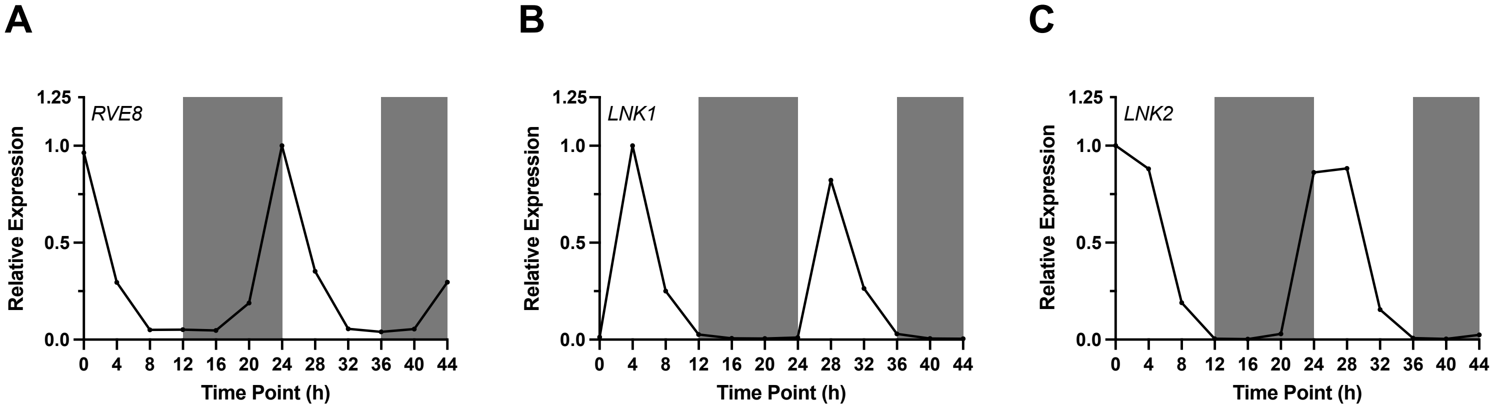


**Supplemental Figure 1 mRNA expression patterns of *RVE8*, *LNK1*, or *LNK2* under photocycles (12 hr light: 12 hr dark).** White and dark grey shading indicates lights-on and lights-off, respectively. Microarray data from diurnal.mocklerlab.com.
