## Supplemental Figure 2 for "COR27/28 Regulate the Evening Transcriptional Activity of the RVE8-LNK1/2 Circadian Complex"

```
TCF1                 1 MNGE--GKLETGGTSAAATPKEEKEEEEVVSQRVVYMWGYLPGASPQRSP     48
                       |||.  ||:..|...|.                ||||.||||||:.::||
RCC1L                1 MNGNIVGKVPIGECKAT----------------VVYMSGYLPGAASEKSP     34

TCF1                49 LMSPVEVKIPPAVE--SSWKDVSGGGCGFAMATAESGKLITWGSTDDLGQ     96
                       ::|||.|::..||.  .|||||.|||||||||.:|.|||||||||||.||
RCC1L               35 ILSPVPVRLSAAVHGGDSWKDVCGGGCGFAMAISEKGKLITWGSTDDEGQ     84

TCF1                97 SYVTSGKHGETPEPFPLPPEVCVQKAEAGWAHCVAVTENQQVYTWGWREC    146
                       |||.||||||||||||||.|..|.:|.:|||||..|||..:.:||||:||
RCC1L               85 SYVASGKHGETPEPFPLPTEAPVVQASSGWAHCAVVTETGEAFTWGWKEC    134

TCF1               147 IPTGRVFGQVDGDSCERNISFSTEQVSSSSQGKKSSGGTSSQVEG-RGGG    195
                       ||:        .|...:..|.|:||||.:|||..::.||:.|.|. :.|.
RCC1L              135 IPS--------KDPVGKQQSGSSEQVSPASQGSNAASGTTLQNENQKVGE    176

TCF1               196 EPTKKRRISPSKQAAENSSQSDNIDLSALPCLVSLAPGVRIVSVAAGGRH    245
                       |..|:||:|.:|...|..:...:. .:..|.|||:..||||.|||.||||
RCC1L              177 ESVKRRRVSTAKDETEGHTSGGDF-FATTPSLVSVGLGVRITSVATGGRH    225

TCF1               246 TLALSDIGQVWGWGYGGEGQLGLGSRVRLVSSPHPIPCIEPSSYGKATS-    294
                       ||||||:||:||||||||||||||||:::|||||.|||:|....||..|
RCC1L              226 TLALSDLGQIWGWGYGGEGQLGLGSRIKMVSSPHLIPCLESIGSGKERSF    275

TCF1               295 ---SGVNMSSVVQCGRVLGSYVKKIACGGRHSAVITDTGALLTFGWGLYG    341
                          .|...::..|..|..|.|:|.|:|||||||.|||.|.|:||||||||
RCC1L              276 ILHQGGTTTTSAQASREPGQYIKAISCGGRHSAAITDAGGLITFGWGLYG    325

TCF1               342 QCGQGSTDDELSPTCVSSLLGIRIEEVAAGLWHTTCASSDGDVYAFGGNQ    391
                       |||.|:|:|:|.|..||.:..:|:|.||||||||.|.||||.||||||||
RCC1L              326 QCGHGNTNDQLRPMAVSEVKSVRMESVAAGLWHTICISSDGKVYAFGGNQ    375

TCF1               392 FGQLGTGCDQAETLPKLLEAPNLENVNVKTISCGARHTA----VITDEGR    437
                       |||||||.|.||.||:||:..|||..:.|.:||||||:|    |:|...
RCC1L              376 FGQLGTGTDHAEILPRLLDGQNLEGKHAKAVSCGARHSAVLAVVLTRRN-    424

TCF1               438 VFCWGWNKYGQLGIGDVIDRNAPAEVRIKDCFPKNIACGWWHTLLLGQPT    487

RCC1L              425 --------------------------------------------------    424

TCF1               488 L    488

RCC1L              425 -    424
```

**Supplemental Figure 2 Protein alignment of TCF1 (AT3G55580) and RCC1L (AT3G53830).** Protein sequences were aligned using the needle algorithm using the EBLOSUM62 matrix, a gap penalty of 10.0, and an extend penalty of 0.5. Sequences share 49.7% identity.
