## Supplemental Figure 3 for "COR27/28 Regulate the Evening Transcriptional Activity of the RVE8-LNK1/2 Circadian Complex"

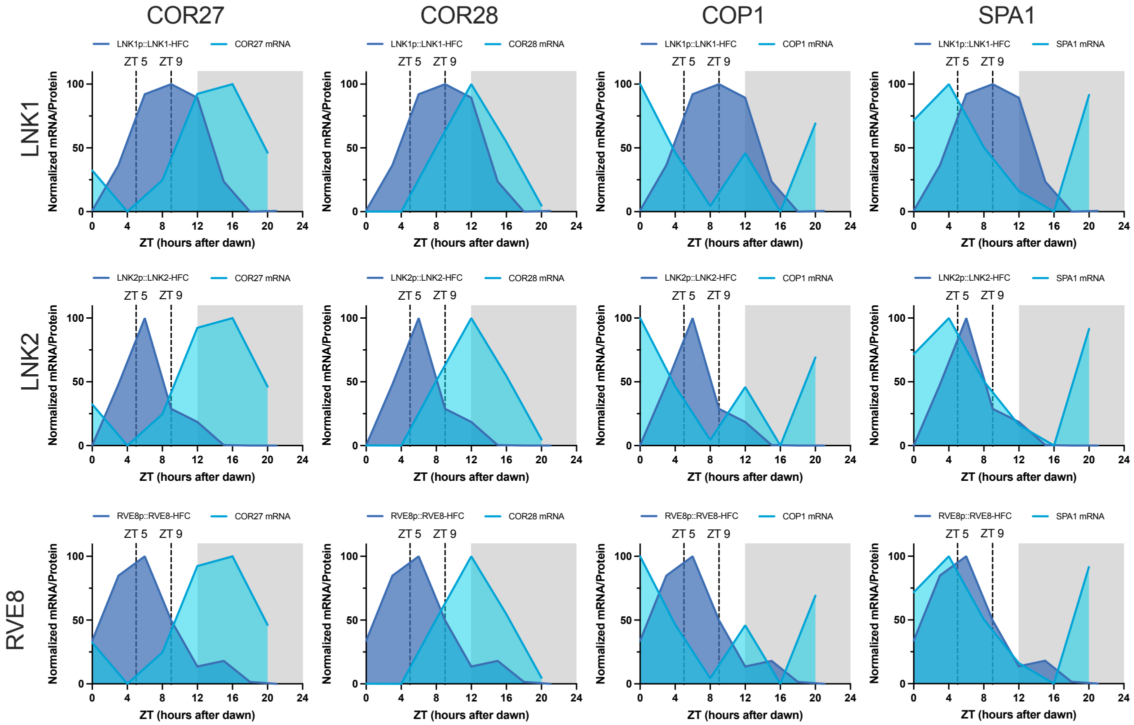


**Supplemental Figure 3 Comparison of HFC-tagged protein abundance with *COR27/28*, *COP1*, and *SPA1* mRNA expression profiles.** 24-hour (12 hr light: 12 hr dark, 22 °C (LDHH)) protein abundance (dark blue) is quantified from Western blots shown in Figure 1D-F. LDHH mRNA data from diurnal.mocklerlab.com (light blue) is overlayed. Vertical dotted lines show the time of day when tissue was collected for APMS. White and grey shading indicated lights-on and lights-off, respectively.
