## Supplemental Figure 4 for "COR27/28 Regulate the Evening Transcriptional Activity of the RVE8-LNK1/2 Circadian Complex"

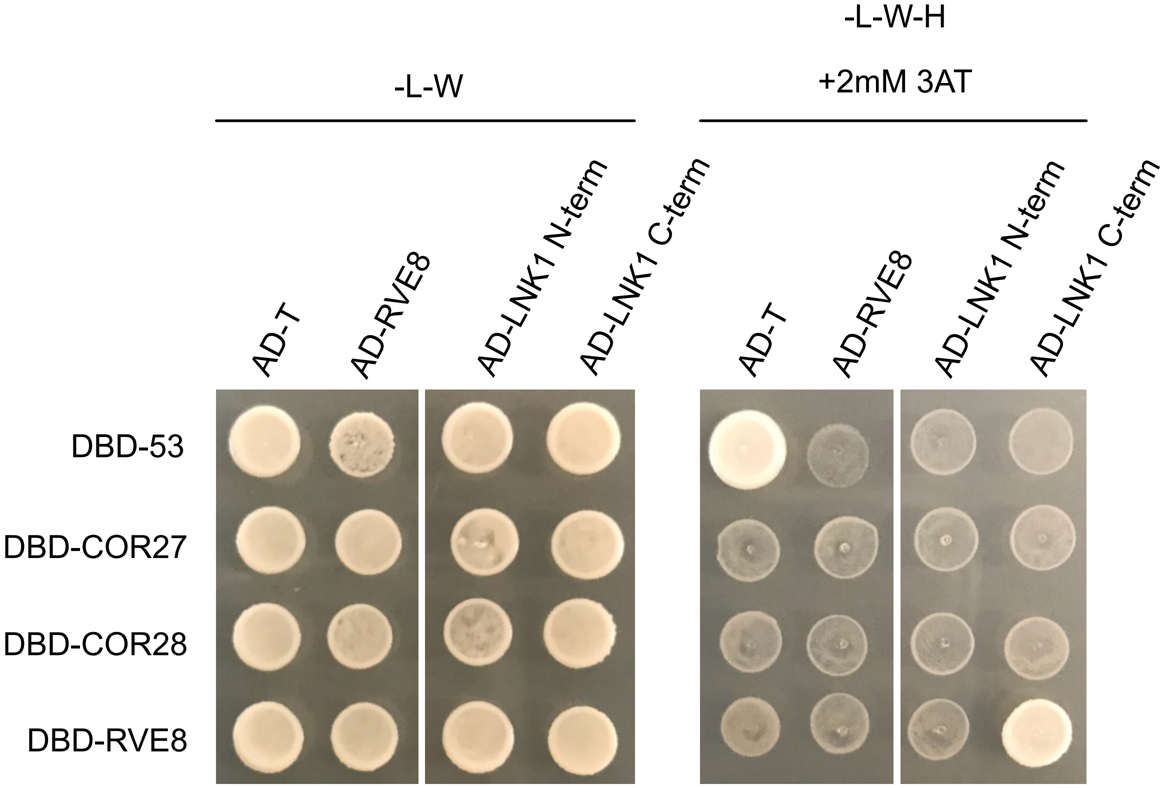


**Supplemental Figure 4 COR27/28 do not interact with RVE8 or LNK1 in a binary Y2H system** Yeast strains Y2H Gold or Y187 expressing pGBKT7 (Gal4-DBD) or pGADT7 (Gal4-AD), respectively, were mated and plated onto selective media. Successful matings were able to grow on -Leucine/-Tryptophan media (-L-W) while positive interactors can grow on -Leucine/-Tryptophan/-Histidine + 2mM 3-amino-1,2,4-triazole (3AT) (-L-W-H +3AT). Only the positive controls DBD-53 (p53) + AD-T (large T-antigen protein) and DBD-RVE8 + AD-LNK1 C-term show an interaction.
