## Supplemental Figure 5 for "COR27/28 Regulate the Evening Transcriptional Activity of the RVE8-LNK1/2 Circadian Complex"

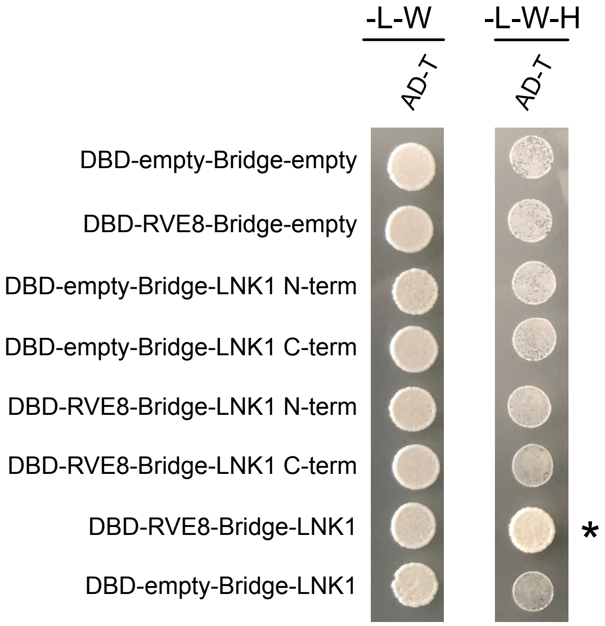


**Supplemental Figure 5 Full-length LNK1 auto-activates in yeast when paired with a DBD-containing protein** Yeast strains Y2H Gold or Y187 expressing pBridge (Gal4-DBD and a Bridge protein) or pGADT7 (Gal4-AD), respectively, were mated and plated onto selective media. Successful matings were able to grow on -Leucine/-Tryptophan media (-L-W). Full length LNK1 (bridge protein, no AD domain) paired with the transcription factor RVE8 (*) can aberrantly activate the expression of the histidine biosynthesis reporter, allowing it to grow on -Leucine/-Tryptophan/-Histidine (-L-W-H) when paired with the negative control large T-antigen protein (T). LNK1 N- and C-terminal truncations do not autoactivate.
