## Supplemental Figure 6 for "COR27/28 Regulate the Evening Transcriptional Activity of the RVE8-LNK1/2 Circadian Complex"

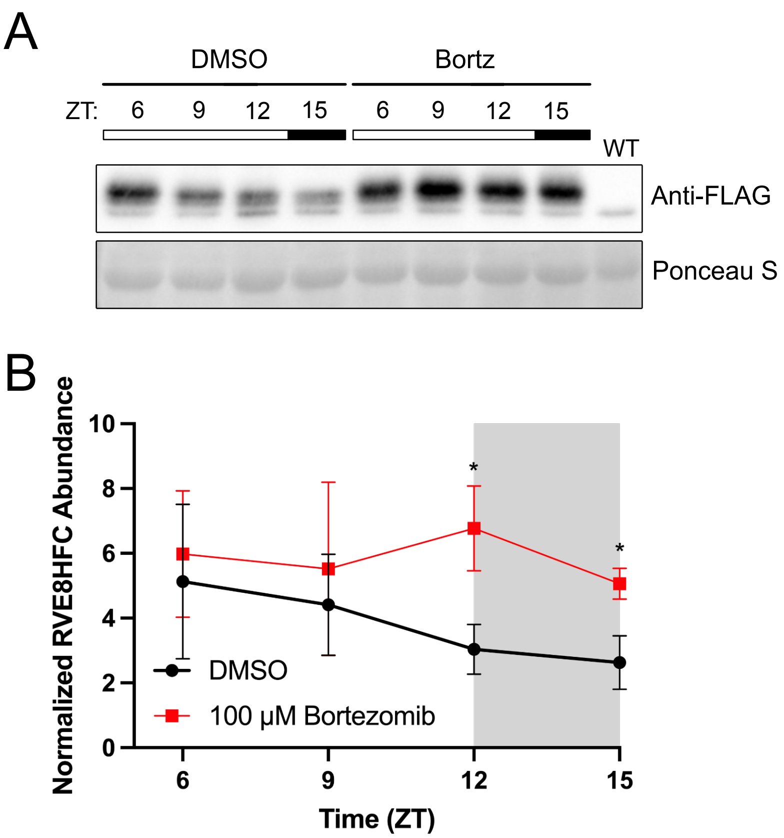


**Supplemental Figure 6 RVE8-HFC protein abundance patterns are regulated by the 26S proteasome** (A) Representative Western blot showing protein expression patterns of RVE8-HFC plants treated with DMSO or 100 µM bortezomib. At ZT5, 12-day-old seedlings growing under 12 hr light: 12 hr dark, 22 °C conditions were immersed in 1/2X MS media containing either 100 µM bortezomib or DMSO. Tissue was collected every 3 hours starting at ZT6. RVE8-HFC was detected with anti-FLAG and Ponceau S staining was used to show loading. (B) Densitometry quantification of RVE8-HFC abundance in (A) normalized to Ponceau S. Points represent the average normalized RVE8-HFC abundance from 3 independent bioreps. Asterisks indicate significant differences between genotypes based on Welch's t-test (* $p<0.05$). Error bars = SD. White and grey shading indicate lights-on and lights-off, respectively. ZT= Zeitgeber Time.
