## Supplemental Figure 7 for "COR27/28 Regulate the Evening Transcriptional Activity of the RVE8-LNK1/2 Circadian Complex"

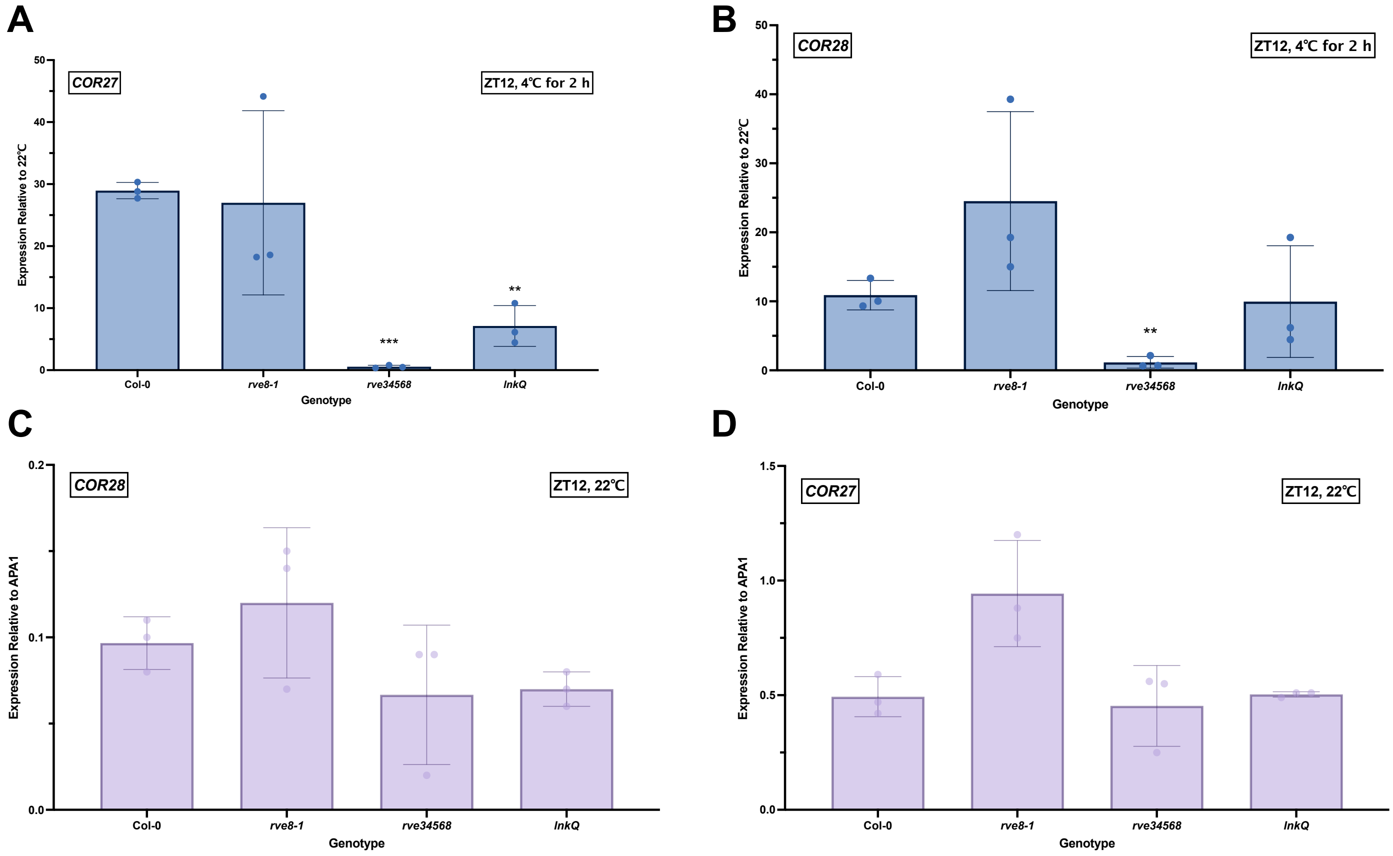


**Supplemental Figure 7 The RVEs and LNKs are important for cold induction of *COR27/28*.** Seedlings were grown on 1/2X MS + 1% sucrose for seven days under 12 hr light; 12 hr dark 22 °C conditions and then transferred at ZT10 to either 22 °C or 4 °C for two hours and tissue was collected at ZT12. (A-B) show the induction of *COR27/28* expression at 4 °C compared to 22 °C . Figures (C-D) show *COR27/28* expression levels at 22 °C . Expression was normalized to the endogenous control gene APA1. Bars show average expression with error bars = SD from 3 independent bioreps (points) for each genotype. Asterisks indicate significant differences as determined by Welch's t-test (** $p<0.01$, *** $P<0.001$).
