## Supplemental Figure 8 for "COR27/28 Regulate the Evening Transcriptional Activity of the RVE8-LNK1/2 Circadian Complex"

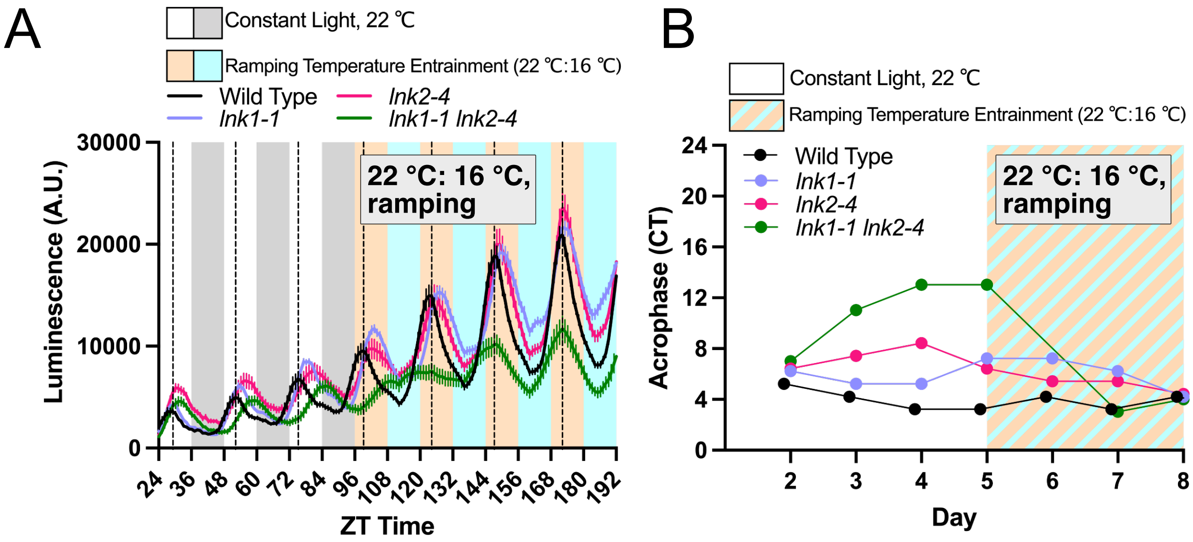


**Supplemental Figure 8 LNK1/2 mutants are also impaired in temperature entrainment under ramping temperature cycles** (A) Luminescence from 7-day-old plants entrained under 12 hr light: 12 hr dark, 22 ℃ conditions expressing a CCA1p::LUC reporter was imaged for at least 3 days in continuous light and temperature (22 ℃) before the chamber was switched to a ramping temperature entrainment program that gradually oscillated between a low temperature of 16 ℃ at ZT16 and a high of 22 ℃ at ZT4. Lines represent the average luminescence from n=16 seedlings with errors bars = SEM. Vertical dotted lines correspond to the peak expression time (acrophase) of the CCA1p::LUC reporter in wild type plants. (B) Acrophase, or time of peak reporter expression, is plotted for each genotype for each day of imaging in constant light and the temperature entrainment condition. Each point represents the acrophase of the averaged luminescence trace shown in (A). CT = Circadian Time. A.U. = Arbitrary Units. ZT= Zeitgeber Time.
